# Dual reinforcement-learning network modules for modeling decision-making with multiple strategies

**DOI:** 10.64898/2026.03.07.709953

**Authors:** Hayato Maeda, Shuo Wang, Akihiro Funamizu

## Abstract

Animals and humans use multiple behavioral strategies to perform tasks. However, neural implementations of multiple strategies remain elusive, as some studies propose distinct pathways, while others observe overlapping brain regions associated with strategies. We propose a hybrid deep reinforcement learning (H-DRL) method, in which one network model implements model-free and inference-based behaviors through synaptic plasticity and recurrent activity. H-DRL uses a single updating rule and switches the strategy according to task demands without an explicit arbitrator. H-DRL reproduced mixed strategies of humans in a two-step task. In the mouse perceptual decision-making task, H-DRL adapted the recurrent dynamics with rich learning when the task condition required inference-based behavior, while adopting model-free behavior with lazy learning for a simple condition. The activity of H-DRL units showed condition-dependent maintenance of previous events, consistent with orbitofrontal cortical activity in mice. Our model provides a unified view that one cortical network automatically determines strategies in use depending on task conditions.

## Introduction

Animals and humans flexibly use, integrate, and switch among multiple behavioral strategies for decision-making. Model-free strategies rely on direct experience whereas model-based strategies use knowledge of state transitions when model-free learning is insufficient^1–3^. Other approaches include inference-based strategies^4,5^ and successor representations^6,7^. Although many studies show that multiple strategies operate within a task, it remains unclear how neural circuits store and implement them^1,3,8,9^. This study proposes a network model that embeds two distinct strategies within a single decision-making network.

Behavioral strategies are examined with tasks such as the two-step decision task, which separates model-free from model-based control^1,3^. Subjects typically use a mixture of both, motivating models with parallel pathways weighted by an arbitrator (**Fig. 1a**) ^1–3^. Some biological evidence supports this division: prefrontal cortex and hippocampus support model-based control^1,10,11^, while ventral striatum and dopaminergic systems support model-free learning via reward prediction errors. Cortico-basal ganglia circuits^12–14^ and OFC/ACC^5^ also show distinct roles for different strategies. However, other studies report overlapping brain regions representing multiple strategies^3,8,15,16^, leaving open how neural networks integrate them to guide choices.

**Figure 1.**
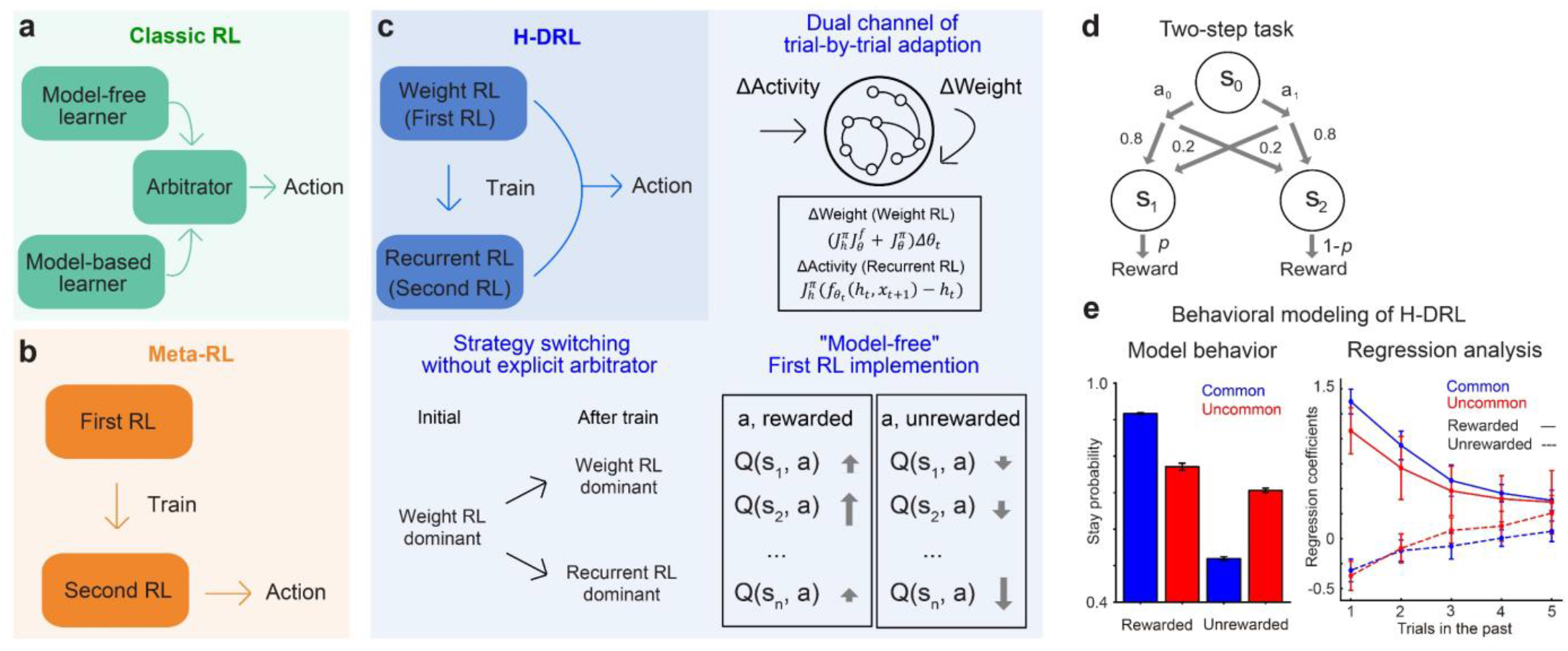
Proposal and validation of hybrid deep reinforcement learning. **a**. Dual-strategy model. Model-free and model-based RLs independently compute reward expectations of choices. The expectations are integrated by an arbitrator. The model-free and model-based RLs use reward- and state-prediction errors, respectively. **b**. Meta-RL. Model-free RL (the first RL) updates synaptic weights of the RNN during the training phase. In the test phase, meta-RL uses the recurrent dynamics (the second RL) to realize adaptive choices. **c**. H-DRL. We modified meta-RL by adding trial-by-trial updates of synaptic weights. H-DRL has three features. (1) Weight updates (weight RL) and recurrent dynamics (recurrent RL) both contribute to sequential choice learning in a single network. (2) The balance between these two RLs is automatically adjusted by the task structure without explicit parameters. (3) Weight RL is implemented as a model-free controller. **d**. Schematic of the two-step task.^3,11^ Actions in the first stage (S_0_) lead to the second stage with fixed transition probabilities (S_1_ or S_2_). Each second stage delivers rewards with certain probabilities that change stochastically during a session. **e**. Output analysis of H-DRL. We trained 20 instances of H-DRL with 100 sessions consisting of 100 trials each. Each instance was initialized with different random weights. The outputs of the model in the last 25 sessions were analyzed. (Left) The probability of the same consecutive choices after common and uncommon state transitions. (Right) Five most recent trials’ regression coefficients for modeling the current choice. Mean ± SD of 20 instances are shown.

Recent years have seen a shift toward deep learning-based approaches. In reward-driven behavior, deep reinforcement learning (deep RL) is commonly employed. A key advantage of deep RL over classical RL is that it enables analysis of the potential neural mechanisms underlying behavior, offering insights into the brain’s complex computational processes^17^. Notably, meta-RL based on the RL^2^ framework has become widely used in neuroscience^18–22^.

A central feature of meta-RL is its ability to autonomously acquire adaptive learning algorithms across diverse tasks^19^. Conventional RL methods are constrained by predefined algorithms, which limit their applicability to tasks that do not conform to their structural assumptions. In meta-RL, however, the predefined RL algorithm (first RL) does not directly compute action values, but instead serves as a learning signal that trains a recurrent neural network (RNN) (second RL). After training, the agent solves tasks solely through the second RL (**Fig. 1b**). By enforcing a strict separation between learning and inference, meta-RL theoretically functions as an approximator of Bayes-optimal solutions^23,24^. In neuroscience, this framework has been proposed as a mechanism by which dopamine trains cortical circuits, particularly within the prefrontal cortex^20^.

As meta-RL has an ability to gradually acquire near-optimal strategies, at the computational level, it can be regarded as a generalization of model-based RL, consistent with a study showing exclusively model-based behavior in the two-step task^20^. However, this advantage in machine-learning side possibly stands in tension with biological evidence showing that animals employ multiple strategies.

Several neuroscience and machine learning studies have proposed additive extensions of meta-RL, such as incorporating an external model-free RL module^25^ or modifying the loss function to incorporate objectives beyond reward maximization^26,27^. These extensions represent possible routes for enhancing meta-RL by increasing architectural complexity.

Instead, we revisit a core assumption of meta-RL and propose the simple modification for implementing multiple behavioral strategies. Although meta-RL includes a predefined first-RL algorithm distinct from the second-RL used for inference, the design principle of timescale separation prevents this first-RL component from influencing behavior. We propose that allowing the predefined RL to serve not only as a learning signal but also as a short-timescale behavioral adaptation mechanism preserves the essential structure of meta-RL while enabling the coexistence of two distinct strategies (**Fig. 1c**). Implementationally, this requires only adopting trial-by-trial online weight updates. With this change, (i) first-RL contributes directly to sequential behavioral adjustments through rapid weight updates, and (ii) the accumulation of these updates shapes the RNN’s recurrent dynamics over the long term, promoting autonomous strategy acquisition as in meta-RL. As a result, behavior emerges from the parallel influences of weight-based updates and recurrent dynamics.

We refer to this framework as hybrid deep reinforcement learning (H-DRL). Although H-DRL differs only minimally from meta-RL in implementation, it embodies a fundamentally different theoretical claim in the context of neuroscience. In H-DRL, trial-by-trial dopamine signals and synaptic plasticity generate a rigid, predefined model-free component (we define weight-RL), whereas the long-term accumulation of dopamine reshapes cortical recurrent dynamics to produce a flexible autonomous RL component (we define recurrent-RL). These two processes co-occur, interact, and potentially compete within a single circuit to generate behavior without an explicit arbitrator.

We observed that H-DRL explained the hybrid use of model-free and model-based strategies in a two-step decision-making task in humans and animals,^3,10^ while the original meta-RL often exhibited purely model-based behavior.^20^ We used H-DRL and modeled the choice behavior of mice in our previous perceptual decision-making task.^15^ The task was performed under repeating and alternating conditions, depending on transition probabilities of sensory stimuli, to distinguish the choice strategies of mice. We observed that H-DRL modeled the condition-dependent choices of mice in terms of choice biases, learning speed, and behavioral strategies.

Perturbation analyses of H-DRL further predicted that weight-RL and recurrent-RL drove the learning of repeating and alternating conditions, respectively. The weight- and recurrent-RLs corresponded to lazy and rich learning of RNNs, and these learning schemes were correlated with the activity of neurons in the mouse OFC. The neurons had the activity-silent mode and the recurrent-dynamics mode for memorizing previous events in the repeating and alternating conditions, respectively. Thus, our proposed H-DRL enabled multiple behavioral strategies with one network implementing lazy and rich learning schemes. Such dual learning engines might be implemented in the OFC network.

## Results

### Proposal and validation of hybrid deep reinforcement learning

The central idea of hybrid deep reinforcement learning (H-DRL) is to remove the strict separation of learning timescales imposed in meta-RL, thereby allowing weight-based adaptation to operate on each trial. Relaxing this constraint enables two fast adaptation processes—weight-based (weight RL) and recurrent-dynamics-based (recurrent RL)—to jointly shape choices within a single recurrent neural network (**Fig. 1c**). Also, for weight-based adaptation to express a model-free component, additional structural conditions beyond trial-by-trial weight updates are required.

To implement this idea, we introduced three key modifications to meta-RL: (i) weight updates were applied on every trial rather than at session boundaries, (ii) updates were computed with simple stochastic gradient descent, and (iii) weights were constrained by softplus nonlinearities to maintain stable online learning. These changes preserve the original architecture while enabling both learning processes to influence behavior on comparable timescales. Other architectural choices and the reasons are described in **Supplementary Note**.

In this framework, H-DRL has three key properties. First, weight RL and recurrent RL coexist within a single RNN. Second, their relative contributions emerge automatically from the task structure rather than being controlled by mixture parameters. Third, trial-by-trial weight updates selectively reinforce rewarded actions and weaken unrewarded ones, consistent with a model-free mechanism (**Fig. 1c**) (**Supplementary Note**).

We tested the performance of H-DRL by maximizing the outcomes during a two-step decision-making task (**Fig. 1d**).^3,10,11^ H-DRL succeeded in simulating the choices of humans and animals, which entailed a hybrid of model-free and model-based strategies (**Fig. 1e**), simply by maximizing the outcomes without fitting the model to the subject data. In contrast, the original meta-RL is known to follow a model-based strategy in which the rewarded choices after common transitions and the unrewarded choices with rare-transitions were equally high probability and the common-unrewarded choices and the rare-rewarded choices were equally low.^20^ These results suggest that H-DRL is able to model choice behaviors using multiple strategies.

### H-DRL explains the choice behavior of mice in the perceptual decision-making task

Next, we used H-DRL and modeled the choice behavior of mice in a perceptual decision-making task in our previous study (**Fig. 2a**).^15^ The task had two rewarded states, and a rewarded state of each trial was controlled probabilistically with transition probabilities (*p*). By setting two different probabilities of repeating (*p* = 0.2) and alternating conditions (*p* = 0.9), our previous study revealed that mice used distinct behavioral strategies.

**Figure 2.**
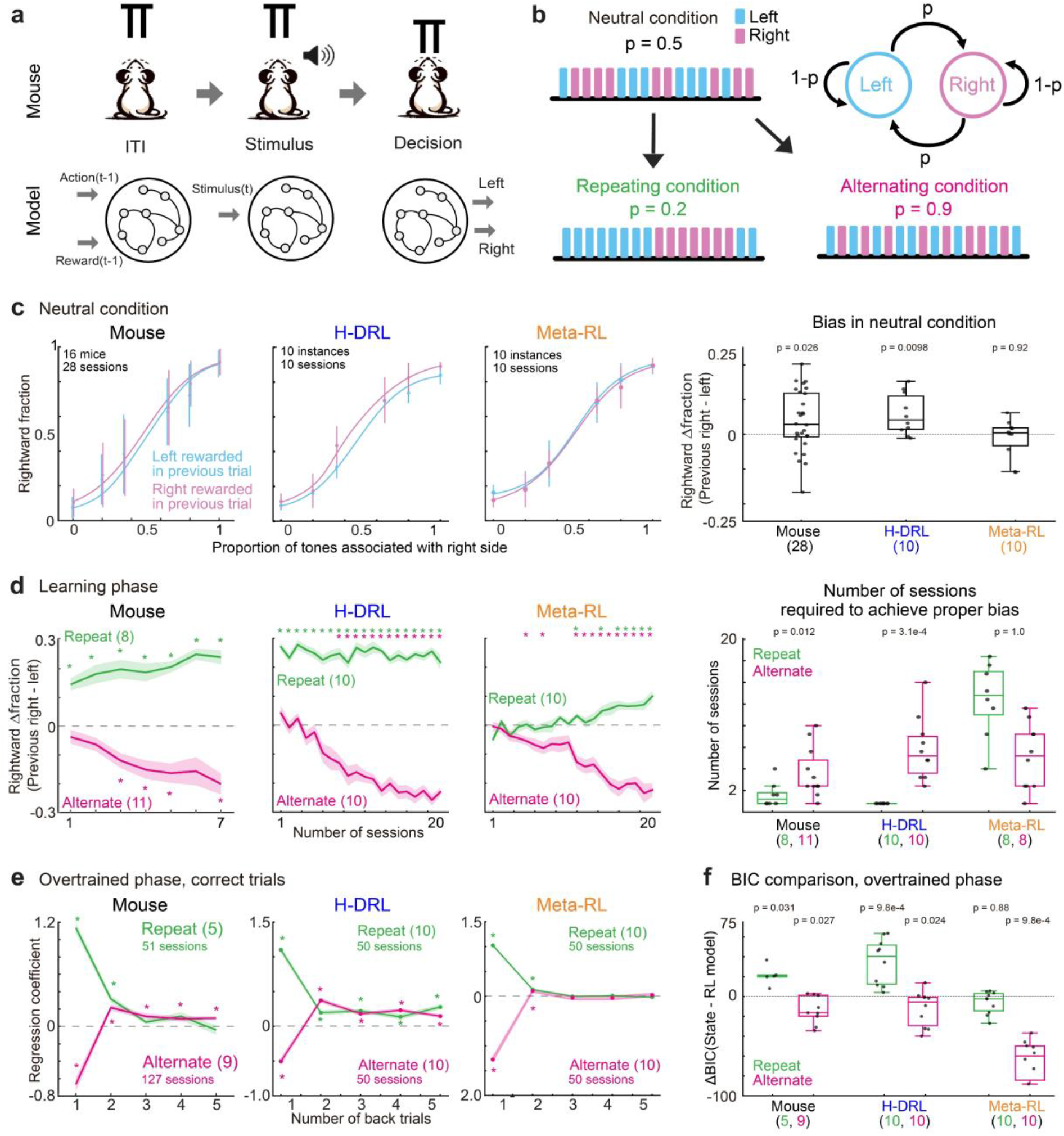
H-DRL models mouse choices with multiple strategies in a perceptual decision-making task. **a**. Schematic of the perceptual decision-making task in our previous study^15^ and model simulation. (Top) Each trial started by moving the spouts away from the mouse. After a random interval of 1–2 s (i.e., ITI), a sound stimulus was presented (Stimulus). The spouts immediately approached the mice after the stimulus ended. The mice licked either the left or the right spout to receive sucrose water (Decision). (Bottom) Each trial in the model was divided into three steps (ITI, Stimulus, and Decision). The action and the reward in the previous trial were used as the inputs during ITI. The stimulus input was applied at the Stimulus step. The output (left or right) of the Decision step was used as the generated choice. **b**. Task conditions. After the mice or the models experienced the neutral condition with a transition probability of *p* = 0.5, they were exposed to either the repeating (*p* = 0.2) or the alternating condition (*p* = 0.9). The original figure was presented in our previous study.^15^ **c-e**. Comparison of choices of mice, H-DRL, and the original meta-RL model. The behavioral results for mice and the original figures are from our previous study.^15^ **c**. Psychometric function in the neutral condition. Mean ± SD are shown. Each model datum was from 10 sessions in each instance with independent initial weights. Right: central mark in the box: median, edges of the box: first quartile (Q1) and third quartile (Q3); bars: most extreme data points without outliers, here and hereafter (a linear mixed-effects model is used for mice; Wilcoxon signed-rank test for models is applied). **d**. Number of sessions required to achieve proper choice biases in the repeating and alternating conditions. The “Rightward Δ fraction” was the difference in the average fraction of right choices after the right- and left-rewarded trials. Mean ± SEM are shown. The mouse data were from 56 sessions of 8 mice in the repeating condition, and 77 sessions of 11 mice in the alternating condition. Data from 200 sessions in 10 instances were used for H-DRL and meta-RL (labels “*” correspond to p < 0.01 in the Wilcoxon signed-rank test). Right: the number of sessions required to bias choices (one-sided Mann–Whitney U test). In meta-RL, two instances did not achieve proper biases in the repeating condition and are not shown in the figure. **e**. Five most recent trials’ regression coefficients for the choices in the current correct trial. Mean ± SEM are shown. Data are represented as five and nine sessions in the repeating and alternating conditions for mice. Data from 50 sessions in 10 instances were used for H-DRL and meta-RL (labels “*” correspond to p < 0.01 in the Wilcoxon signed-rank test). **f**. Model fitting. A model-free strategy and an inference-based strategy were fit to the choices of mice and those generated from the models. For mouse choices, the model fitting and original figures are from our previous study.^15^ The choices generated by H-DRL showed condition-dependent strategies (one-sided Wilcoxon signed-rank test).

To summarize the trial setting, mice were head-fixed and presented with a mixed sequence of low- or high-frequency short pure-tone pulses, called a tone cloud for 0.6 s (stimulus)^28–34^ After the tone, the mice chose the left or the right spout to receive 2.4 *μ*L of 10% sucrose water, depending on the dominant category of tone frequency (i.e., low or high) (decision). Failing to make the proper choice resulted in the mouse being presented with a noise burst for 0.2 s. After an inter-trial interval (ITI) of at least 3.5 s, the next trial started.

In the previous study, the mice were initially trained in the neutral condition in which high- or low-category tones were presented randomly (**Fig. 2b**).^15^ The task then separated the mice into two groups and assigned them to either the repeating or the alternating condition. During the learning phase, the repeating condition had same tone category in every trial with 80 % (*p* = 0.2), while the alternating condition switched the tone category with 90 % (*p* = 0.9). In the overtrained phase, we set the transition probabilities of repeating and alternating conditions at *p* = 0.2 and 0.8, respectively.

In theory, the optimal strategy for both repeating and alternating conditions is an inference-based strategy that estimates the transition probability of tone category between trials. An alternative approach in the repeating condition is a simple model-free strategy that repeats the rewarded choice. In contrast, such a simple model-free strategy in the alternating condition makes it difficult to switch rewarded choices without having state transitions or task-specific state definitions.

We trained our H-DRL model and the original meta-RL model with a simulated perceptual decision-making task with similar trial and task schedules for mice. Each session consisted of 500 trials, and recurrent units’ activity was reset to zero at every session start. Each trial was categorized with respect to three steps (i.e., ITI, the stimulus and the decision) (**Fig. 2a**). During the ITI and the stimulus step, the model received inputs of the previous choice and reward, and a scalar proportion of high-frequency tone in the tone cloud, respectively. During the decision step, the models generated a categorical choice (left or right) by applying a softmax function to the actor output (Meta-RL) or the action-value output (H-DRL) (**Methods**). A binary reward (1 if correct, and 0 otherwise) was provided immediately after the choice.

We first trained the models with a simulated neutral condition until they reached the correct rate of 80%. The H-DRL and meta-RL models took 18.3 ± 8.0 sessions and 128.8 ± 15.5 sessions (mean ± SD from 10 instances), respectively. The same trained weights were used for both the simulated repeating condition and the alternating condition (20 sessions for the learning phase and 5 sessions for the overtrained phase for each condition). Models with 10 independent initial weights (described as 10 instances) were used for the simulation of both neutral-repeating and neutral-alternating conditions. Each simulated session had 500 trials. In the neutral condition, we analyzed the model-generated choices in the last session. For the overtrained phase of repeating and alternating conditions, we analyzed the generated choices in all five sessions. Note that the recurrent connections were not fixed even during the overtrained phase.

We compared the choices generated by the H-DRL and meta-RL models with those of mice already reported in our previous study.^15^ For the mice, (1) the choices were biased depending on the previously rewarded side in the neutral condition (**Fig. 2c**). (2) The acquisition of proper choice biases was faster in the repeating condition than in the alternating condition (**Fig. 2d**)^15^. (3) In the overtrained phase, the last five trials affected the choice in the current trial (**Fig. 2e**). These behavioral observations of the mice were captured by the choices generated by H-DRL but not by the original meta-RL model.

For our perceptual decision-making task, we previously reported that the choices of mice in the repeating and alternating conditions fit model-free and inference-based strategies, respectively^15^ (**Fig. 2f, left**). We therefore fit the choices generated by H-DRL and meta-RL with the two behavioral strategies (**Methods**) (**Fig. 2f**). We observed that the strategy of H-DRL choices was consistent with that of mice, while the strategy of meta-RL was inference-based in both repeating and alternating conditions. Overall, these results suggest that H-DRL has an advantage in modeling mouse choices using multiple strategies in a perceptual decision-making task.

### Perturbation of H-DRL shows the use of weight- and recurrent-RLs for choices in repeating and alternating conditions

In our H-DRL model, the weight RL updated the expected reward of choices (i.e., action values) by changing the connection weights in each trial (**Fig. 3a**). Specifically, weights were updated so that rewarded network states were strengthened in a model-free manner. In contrast, the recurrent RL updated the action values with recurrent dynamics. In our perceptual decision-making task for mice (**Fig. 2a**), proper choice biases in the repeating condition required only a model-free update of action values, while those in the alternating condition were difficult to achieve by a simple updating rule. We thus hypothesized that H-DRL generated the choices in repeating and alternating conditions with the weight- and recurrent-RLs, respectively.

**Figure 3.**
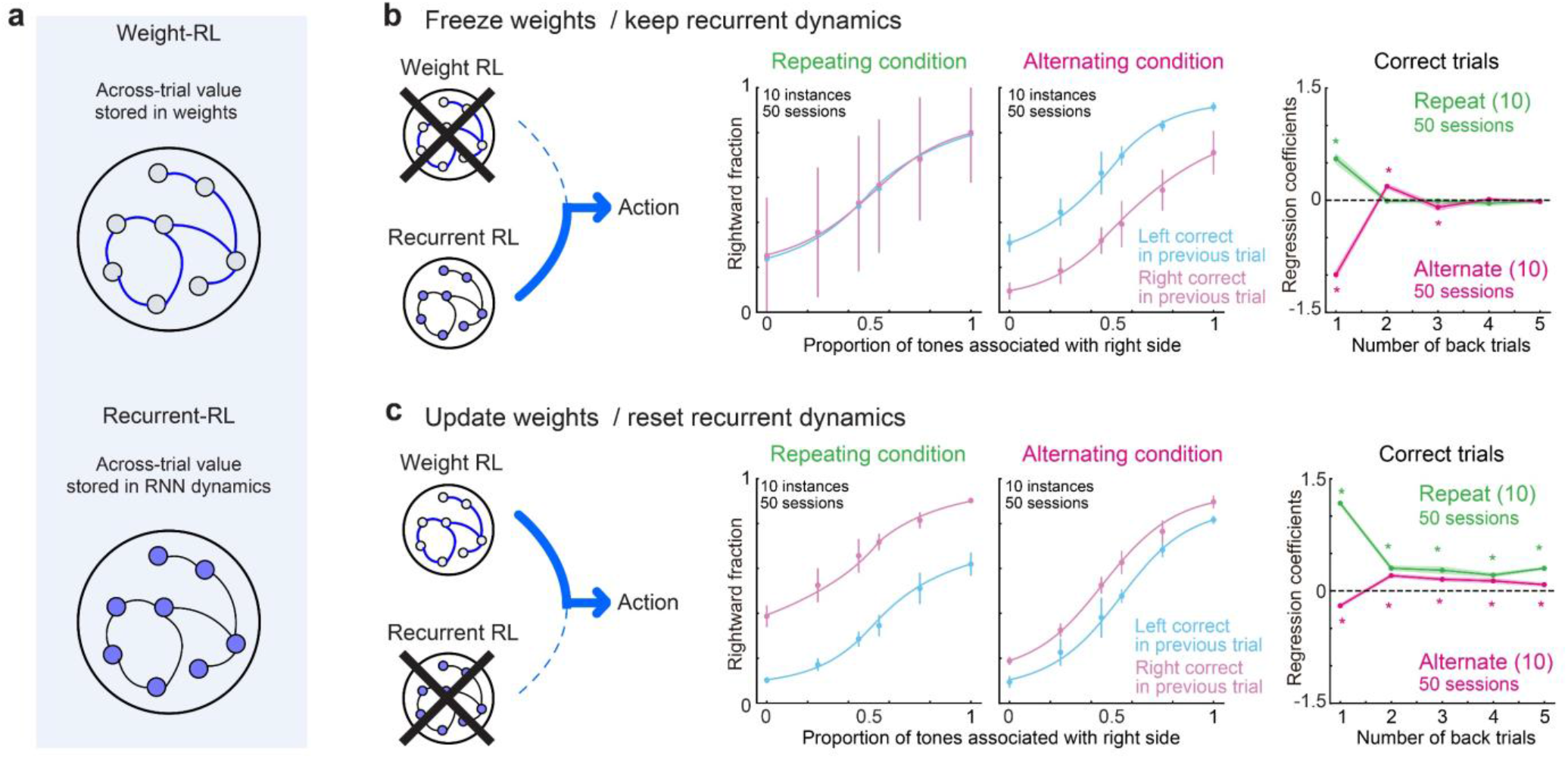
H-DRL automatically changed the RL module in use based on task conditions. **a**. Scheme of weight- and recurrent-RLs that update the action values by changing connection weights and via recurrent dynamics, respectively. **b**. Weight-freeze test. We stopped the updates of connection weights but retained the recurrent dynamics. Middle: psychometric functions under repeating and alternating conditions. Mean ± SD are shown. Right: five most recent trials’ regression coefficients for correct choices. Mean ± SEM are shown (labels “*” correspond to p < 0.01 in the Wilcoxon signed-rank test). Each panel consisted of data from five sessions in 10 independent instances of H-DRL. **c**. Activity-reset test. We updated connection weights with weight RL but reset the activity of all units to zero in each trial. Data presentation is consistent with that in **b**.

To test this hypothesis, we performed two perturbation tests on H-DRL (**Fig. 3b, c**). One was a weight-freeze test that fixed the recurrent-connection weights, similarly to the conventional meta-RL, to stop the use of the weight RL for action learning. The other was an activity-reset test, which reset the activity of all RNN units to zero between the ITI and Stimulus steps in each trial. This activity-reset test prevented the trial-by-trial accumulation of activity within the RNN to remove the recurrent RL.

In the weight-freeze test (**Fig. 3b**), the proper choice biases decreased mainly in the repeating condition but not in the alternating condition. In contrast, the activity-reset test disrupted the proper biases in the alternating condition (**Fig. 3c**). These results suggest that the choice biases in the repeating and alternating conditions in H-DRL were achieved by the weight- and recurrent-RLs, respectively. Such selection of strategies was automatically performed by H-DRL depending on task conditions.

### Recurrent dynamics of H-DRL predict distinct network learning modes in the repeating and alternating conditions

How did H-DRL automatically select the learning modules for repeating and alternating conditions? It is known that an RNN generally changes connection weights to behave as a nonlinear state approximator in a partially observed Markov decision process (POMDP), such as in our perceptual decision-making task.^15,35^ In contrast, if the task can be performed without adjusting the internal representations, the RNN mainly adjusts the output, similarly to reservoir computing. The former and latter approaches correspond to rich and lazy learning, as they need and need not update recurrent dynamics, respectively.^36^

In our perceptual decision-making task (**Fig. 2a**),^15^ H-DRL initially learned the task structure in the neutral condition and generated tone frequency-dependent choices (**Fig. 2c**). As the choices in the neutral condition were already biased by previous events, H-DRL did not need to estimate the transition of rewarded states in the repeating condition. In contrast, the alternating condition required estimating state transitions to reverse the biased behavior.^15,37^ We therefore hypothesized that the RNN performed lazy and rich learning in the repeating and alternating conditions, respectively (**Fig. 4a**).

**Figure 4.**
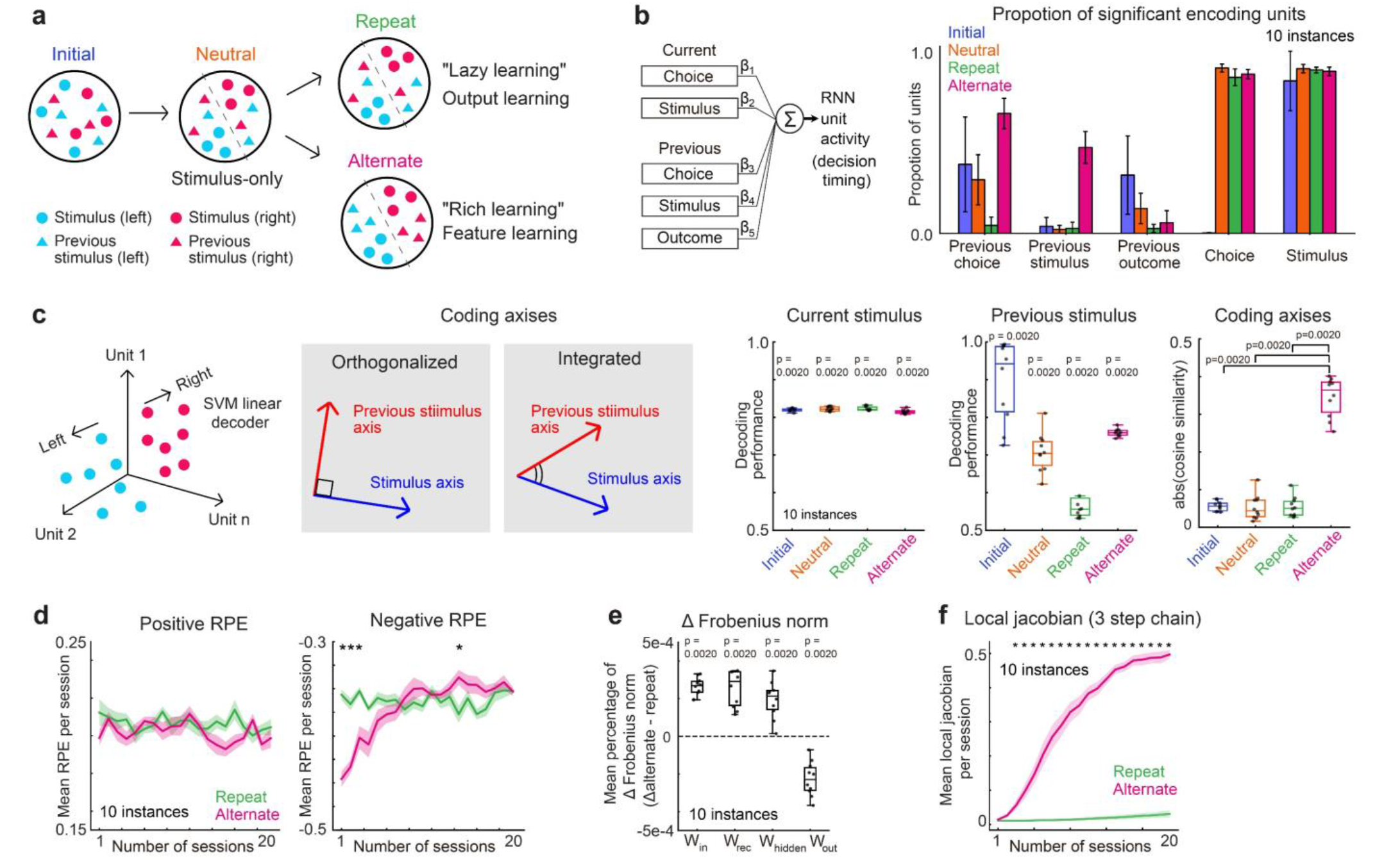
Distinct recurrent dynamics of H-DRL in repeating and alternating conditions. **a**. Hypothesis of how H-DRL automatically selects the learning modules for repeating and alternating conditions. In the neutral condition, H-DRL changes the RNN dynamics from the initial network to learn the stimulus–reward association in the task. In the repeating condition, the RNN keeps the network dynamics (lazy learning), while in the alternating condition, the RNN develops condition-adaptive dynamics (rich learning). **b**. Encoding analysis with a generalized linear model (GLM) with 10-fold cross validation. We analyzed the proportion of units representing the stimuli, choices, and outcomes in the previous and current trials. Mean ± SD are shown. **c**. Population decoding analysis. A support vector machine was used to identify the axis in population neural activity that separated binary task variables (left). Scheme of task-adaptive representation: orthogonal axes indicate independent representations of current and previous stimuli, whereas integrated axes show that the network links the two variables (middle). Decoding performance and cosine similarity of coding axes at four time points (right) (Wilcoxon signed-rank test). **d**. Trajectory of trial-averaged reward prediction errors (RPE). Positive and negative RPEs are plotted separately in the repeating and alternating conditions. Mean ± SD are shown (labels “*” correspond to p < 0.01 in the Wilcoxon signed-rank test). **e**. Average absolute change in weights (Δ Frobenius norm) across trials in each session. The normalized Frobenius norm was used to quantify relative weight updates for each of the four connections in the model (Wilcoxon signed-rank test). **f**. Trial-averaged local Jacobian (spectral radius) averaged across sessions. The local Jacobian represents how recurrent-activity changes propagate over time. Mean ± SD are shown (labels “*” correspond to p < 0.01 in the Wilcoxon signed-rank test).

To test this hypothesis, we analyzed and compared RNN activity dynamics at four different time points: (1) in the initial no-learning phase (Initial), (2) after the neutral condition (Neutral), (3) after the repeating condition (Repeat), and (4) after the alternating condition (Alternate) (**Fig. 4a**). At each time point, to examine the recurrent dynamics of the RNN independently from the weight RL, we froze the weights of H-DRL. All models were used in five sessions (with each session consisting of 500 trials) with a transition probability of 0.5 to prevent potential biases from conditional variabilities.

We first performed an encoding analysis of RNN units’ activity (**Fig. 4b**). A generalized linear model (GLM) was used to investigate whether the activity of RNN units at the Decision step (**Fig. 2a**) represented choice, stimulus, the previous choice, the previous stimulus, and the previous outcome (i.e., five variables and a constant term). Initially, we observed that 48.3 ± 8.9 % (Mean ± SD of 10 instances) of units encoded the previous stimulus in the alternating condition. The representation of the previous stimulus required integrating the RNN inputs of the previous action and outcome, and it did not increase in the neutral (2.2 ± 2.3%) and repeating conditions (2.8 ± 3.4%) compared to the initial random network (3.9 ± 5.0%) (Wilcoxon signed rank test, neutral: p = 0.88; repeat: p = 0.79). Thus, the encoding of the previous stimulus increased selectively in the alternating condition (p = 0.0020 in all three comparisons), consistently with our hypothesis that network dynamics changed in the alternating condition. Notably, the choice-encoding units appeared only after we trained the network in the neutral condition.

Next, we performed a decoding analysis with a linear support vector machine (SVM) (**Fig. 4c**).^38^ The SVM decoded the categorized stimulus of the previous and current trials from the activity of RNN units at the Decision step. As expected, the RNN units’ activity could be used to decode the previous and current stimuli at all four time points (Wilcoxon signed-rank test, p = 0.0020 in all eight cases).

One important aspect of an RNN is that its random connections, even at the initial time point, have linearly separable inputs. In such random connections, separating hyperplanes for different task variables are unrelated, and cosine similarities of hyperplanes are low. In contrast, if recurrent connections are organized by learned task structures, cosine similarities become high. We noted that cosine similarities between the current- and previous-stimulus hyperplanes were greater after the alternating condition than after the other conditions (Wilcoxon signed-rank test, p = 0.0020 in all three comparisons) (**Fig. 4c right**).

Such differences between repeating and alternating conditions might depend on how much the recurrent connections changed with RPEs. We observed that during the initial learning of task conditions, negative RPEs were significantly greater in the alternating condition than in the repeating condition (**Fig. 4d**), possibly because the proper choices in the alternating condition required reversing the biased behavior in the neutral condition by updating recurrent dynamics.

Next, we elucidated how negative RPEs affected network connections by separately analyzing the Frobenius norm of weights in (1) RNN inputs (*W*_*in*_), (2) recurrent connections (*W*_*rec*_), (3) RNN to linear layer (*W*_*hidden*_), and (4) outputs (*W*_*out*_) (**Fig. 4e**). The Frobenius norm was the square root of the sum of squared weight changes across all components in each weight matrix. When we compared weight changes in the repeating and alternating conditions, *W*_*out*_ showed large changes in the repeating condition, while the other weights changed in the alternating condition.

Furthermore, we introduced a local chain Jacobian to investigate how recurrent dynamics facilitated learning through gradient amplification over time (**Fig. 4f**) (**Methods**).^39,40^ The trial-by-trial spectral radius of the local chain Jacobian increased in the alternating condition compared to the repeating condition, suggesting that the internal dynamics of the RNN were affected by RPEs in the alternating condition. Overall, these results suggest that when H-DRL could not maximize the reward in the alternating task, it changed the recurrent connections with recurrent RL following rich learning, whereas it used lazy learning to implement a simple choice strategy in the repeating condition.

### The decoding performance of neurons in the mouse orbitofrontal cortex (OFC) is consistent with H-DRL

Finally, we compared the RNN unit activity of H-DRL with the neuronal activity in mice in our perceptual decision-making task.^15^ H-DRL predicts that the choices in repeating and alternating conditions depend on weight updates (weight RL) and recurrent dynamics (recurrent RL), respectively (**Figs. 2 and 3**). Hence, neurons do not need to memorize previous events with the activity during the repeating condition but need to do so with changes in synaptic connections (e.g., activity-silent working memory).^41–44^ In contrast, neurons in the alternating condition rely on recurrent activity during inter-trial intervals (**Fig. 5a**).

**Figure 5.**
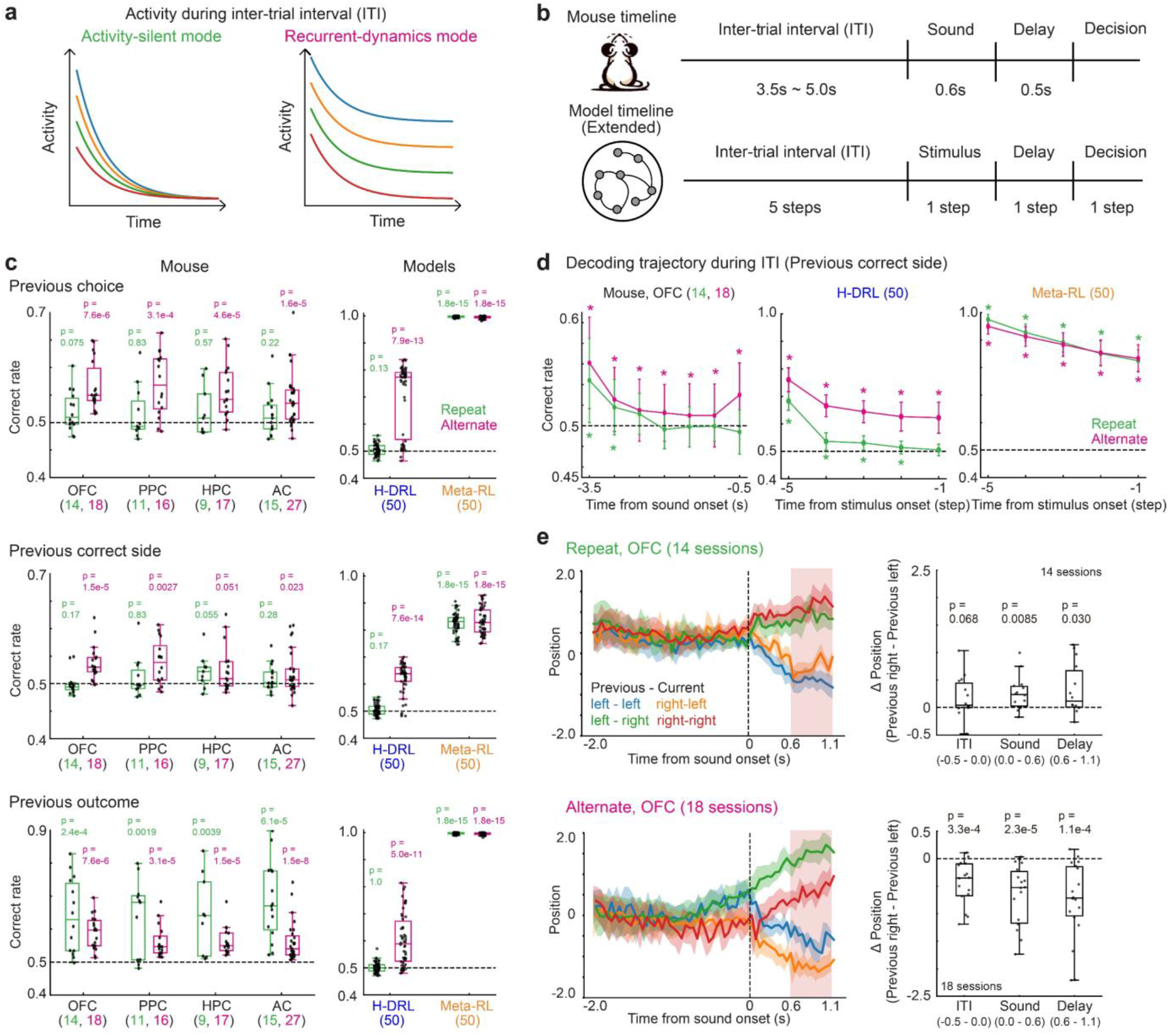
Condition-dependent representations of previous events in the mouse orbitofrontal cortex. **a**. Two modes of maintaining previous events during the inter-trial interval. (Left) The activity-silent mode does not maintain activity during ITI but maintains previous events with synaptic changes linked to the weight RL. (Right) The recurrent-dynamics mode maintains previous events with activity during the ITI, linked to the recurrent RL. **b**. Trial structure of the mouse behavioral task and the H-DRL with extended ITI. Each trial in the mouse task consisted of a random ITI (3.5–5.0 s), the sound stimulus (0.6 s), and a delay (0.5 s) before the spout approached to allow making a choice. The extended H-DRL had five ITI steps, one Stimulus step, one Delay step, and one Decision step. **c**. Performance of decoding previous events from the activity of mouse neurons (left) and model units (right). To prevent crosstalk across the decoding of the previous choice, the outcome, and the correct side, trials were subsampled so that the trial combinations of previous choices and outcomes were equally included in the decoder training. The numbers of sessions for mice and models are shown in parentheses (Wilcoxon signed-rank test). **d**. Decoding of the previous correct side during ITI with the mouse OFC, H-DRL, and meta-RL. Mean ± SD are shown (labels “*” correspond to p < 0.05 in the Wilcoxon signed-rank test). **e**. Population activity of OFC neurons projected to a choice axis during the delay period. (Left) Population neural trajectories categorized with the previous- and current-correct sides. Mean ± SEM are shown. (Right) Differences in population trajectories between trials with previous correct right and left sides. The time from the onset of sound (s) is shown in parentheses (Wilcoxon signed-rank test).

To explore and compare the modes of memorizing past events in mouse neurons and H-DRL units, we analyzed the temporal activity changes during ITIs. In this analysis, we extended the time steps of H-DRL and the original meta-RL, with five time steps of ITI, one Stimulus step, one Delay step, and one Decision step (with eight time steps per trial) (**Fig. 5b**). We verified that such extensions did not alter the choices generated from H-DRL (**Fig. S1**).

For mouse neural activity, we used the data from the OFC, the posterior parietal cortex (PPC), the hippocampus (HPC), and the auditory cortex (AC), where neural activity was recorded during both the repeating and alternating conditions. We used the sessions for analysis when more than 50 neurons were recorded simultaneously (**Methods**).

During the late timing of ITI (i.e., from −0.5 s to 0 s relative to the onset of sound), we investigated the decoding performance of mouse neural activity for the previous choice, the previous correct side and the previous outcome with a sparse logistic regression (SLR) (**Fig. 5c**). To prevent a potential bias from the number of neurons, we randomly selected 50 neurons for each decoding and performed a 10-fold nested cross validation.^29^ We ran the decoding analysis 100 times and analyzed the average performance in every session. For H-DRL and meta-RL, we used all 100 units’ activity during the last step of ITI and investigated the performance of decoding previous events. The observed performance was validated with SLR, following the same approach as in the case of mouse neural activity.

In mouse neurons, although the performance of decoding the previous outcome was beyond the chance level in both the repeating and alternating conditions, decoding of the previous choice and the correct side was achieved only in the alternating condition, except for HPC in the case of decoding the previous correct side (**Fig. 5c**). These differences across task conditions were consistent in the unit activity of H-DRL but not in that of meta-RL. For H-DRL, the performance of decoding previous events was beyond chance levels only in the alternating condition. In contrast, the performance for meta-RL was consistent across task conditions. Although there was inconsistency between the activity of mice and H-DRL in the decoding of previous outcomes, the decoding performance of H-DRL was similar to that of mouse neurons, which was not the case in the original meta-RL.

Next, we examined how the performance of decoding previous events was temporally modulated during ITI (**Fig. 5d**).^43^ For mouse neurons, we considered performance every 0.5 s between −3.5 and 0 s relative to the onset of sound. In OFC, the performance of decoding the previous correct side decreased gradually, especially in the repeating condition, and approached the chance level at late timings of ITI. Similarly, the decoding performance of H-DRL was significantly above the chance level in the beginning of ITI in both conditions, and decreased to the chance level in the repeating condition at late time steps. In contrast, in meta-RL, decoding performance deteriorated gradually, but its measures were similar between the alternating and repeating conditions. In PPC, HPC, and AC (**Fig. S2**), the performance of decoding previous events differed from that in OFC. The performance of decoding the previous correct side in PPC was high only in the alternating condition, while it was maintained in neither condition in HPC or AC.

Finally, we investigated how the neural representations of previous events were maintained during the current trial (**Fig. 5e, Fig. S3**). During the delay period (i.e., 0.6–1.1 s from the onset of sound), we decoded the current choice with SLR. We then used the choice-decoding axis from SLR to project the neural activity between −2.0 s and 1.1 s from the onset of sound (**Fig. 5e, left**). In the OFC, in the repeating condition, previous events affected choice activity during the sound and delay periods, whereas in the alternating condition, they also affected choice activity during ITI (**Fig. 5e, right**). In PPC, HPC, and AC, choice-dependent neural separations were modulated by previous events only in the alternating condition (**Fig. S3**). These results again suggest the difference in neural maintenance of previous events between task conditions.

## Discussion

This study proposed the H-DRL model that implemented multiple behavioral strategies with dual RL modules in one network (**Fig. 1**). H-DRL makes three major changes to a conventional meta-RL^20^ to model choices of humans and animals (**Fig. 1c**). (1) It has dual learning modules of first and second RLs (defined as weight- and recurrent-RLs) for both action learning and execution in each trial. (2) The dependencies of the two RLs for choices are adjusted in each task without explicit parameters. (3) H-DRL relies on online updates of recurrent weights in every trial to strengthen the previous rewarded choice and weaken the previous unrewarded choice in a model-free manner.

H-DRL succeeded in simulating a mix of model-free and model-based strategies in a two-step decision-making task^3,10,11^ (**Fig. 1d, e**). H-DRL also modeled mouse choices in our perceptual decision-making task, which required mice to use distinct strategies in repeating and alternating conditions (**Fig. 2**).^15^

A perturbation of H-DRL showed that H-DRL automatically selected the RL module in use depending on task conditions (**Fig. 3**). In the repeating condition, the weight RL mainly changed the connection weights of the outputs (**Fig. 4e**), suggesting lazy learning in the RNN. In contrast, in the alternating condition, the recurrent dynamics were updated with negative RPEs to represent previous stimuli, suggesting rich learning (**Fig. 4b-f**).^15,35^

Finally, we observed that such differences in learning modules in repeating and alternating conditions were consistent with the neuronal activity in mice, especially in OFC (**Fig. 5**). During ITI in the repeating and alternating conditions, the neuronal activity did not (respectively, did) represent the previous choice and the correct side. As mice biased the choices depending on the previous events in the task, these results imply that different network modes (i.e., the activity-silent mode and the recurrent-dynamics mode) maintain the task history in OFC, possibly consistently with lazy and rich learning of the RNN (**Fig. 4**).

Proper choice behavior requires memorizing past events. Early physiological^45–47^ and modeling studies^48,49^ of working memory (WM) revealed that recurrent dynamics of RNN memorize stimulus features with persistent neural activity. An RNN also models a sequence of population activities for WM in mice.^50,51^ In contrast to using recurrent dynamics, recent studies propose an activity-silent mode that utilizes short-term synaptic plasticity for memorizing past events.^41–44^ As both recurrent dynamics and synaptic plasticity are important for flexible behavior, our H-DRL model has biologically-inspired dual learning modules of recurrent- and weight-RLs to maintain past events in one network.^52^

Consistently with previous studies in humans and animals,^1–3,8,11^ dual learning modules in H-DRL were essential for modeling mixed behavioral strategies (**Fig. 2**). Our H-DRL analysis further proposed that lazy and rich learning in the RNN corresponded to the model-free and adaptive inference-based strategies, respectively.^36^ Earlier studies have shown that the frontal cortex serves as a computing reservoir using a high-dimensional recurrent network to perform a task similarly to lazy learning^53,54^ while also attaining a task-dependent low-dimensional structure by changing recurrent dynamics through rich learning.^55^ Our study unifies such past research and proposes that learning schemes of a recurrent cortical network are task-dependent (**Fig. 3**).

The OFC is known to represent both model-free and inference-based strategies.^5,56,57^ Synaptic plasticity and neural activity in the OFC, corresponding to the weight- and recurrent-RLs in this study, are necessary for session-by-session and trial-by-trial learning of tasks, respectively.^21^ Although we mainly observe a correlation between the activity of the OFC and H-DRL, other brain regions are also involved in multiple behavioral strategies.^11,58,59^ It is thus essential to explore how other regions are related to H-DRL.

H-DRL has several limitations. It uses truncated backpropagation through time (TBPTT), which is limited by the number of steps for updating weights (**Methods**).^60^ A possible improvement of H-DRL is to use real-time recurrent learning (RTRL) weight updates.^61^ Additionally, further animal experiments are necessary for testing the predictions from H-DRL (**Figs. 3, 4**). First, this study did not include perturbation experiments in mice. Perturbations and measurements of both synaptic plasticity and neural activity are essential for validating the role of weight- and recurrent-RLs in decision–making.^21,62,63^ Second, it would be beneficial to record and analyze the neuronal activity of mice while learning to investigate whether the changes in network dynamics are consistent with the H-DRL prediction (**Fig. 4**). In summary, our proposed H-DRL approach provides a network model of how animals make choices using multiple behavioral strategies.

## Methods

### Model Architectures

#### Meta-RL model

We used widely adopted meta-RL frameworks.^18–20^ The specific framework implemented a deep reinforcement learning model combining recurrent neural networks and RL algorithms. The RNNs in this study consisted of 100 fully connected long short-term memory (LSTM) units.^64^ At each time step *t*, the network received an observation vector *x*_*t*_ ∈ ℝ^D^. LSTM units were composed of an input gate, a forget gate and an output gate, each modulated by a learned function. The dynamics of the LSTM unit were defined as follows:

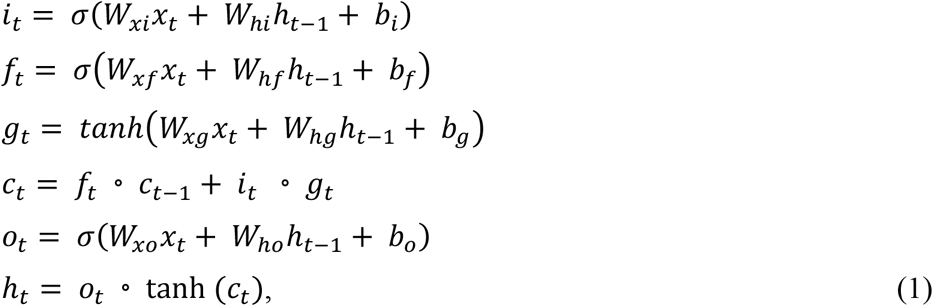

where *h*_*t*_, *c*_*t*_ ∈ ℝ^100^, *W*_*x*\*_∈ ℝ^100×D^,*W*_*h*\*_ ∈ ℝ^100×100^, and biases *b*_*_ ∈ ℝ^100^. Both the policy and the state value (the expected reward of the current state) were read out from the same hidden activations *h*_*t*_ ∈ ℝ^100^:

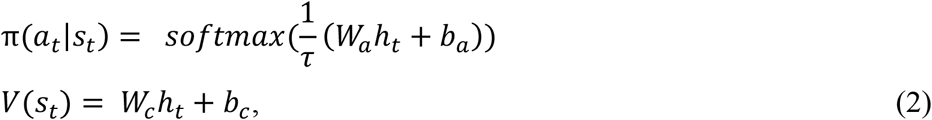

where *W*_*a*_ ∈ ℝ^2×100^, *W*_*c*_ ∈ ℝ^1×100^, *b*_*a*_ ∈ ℝ^2^, and *b*_*c*_ ∈ ℝ. *τ* was a temperature hyperparameter that controlled policy entropy (set to 0.2). Action *a*_*t*_ was sampled from *π*(*a*_*t*_|*s*_*t*_).

We trained the meta-RL model using an Advantage Actor-Critic (A2C) algorithm,^65^ with a truncated backpropagation-through-time (TBPTT) and root-mean-square-propagation (RMSProp) (alpha = 0.99). Following the A2C method, we defined the n-step return and advantage as follows:

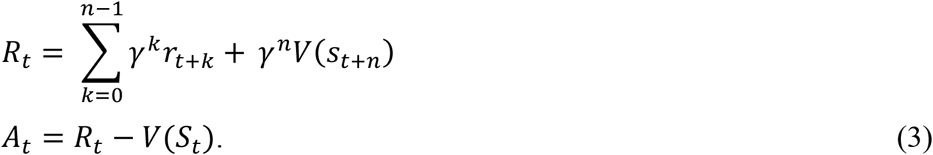

The losses were

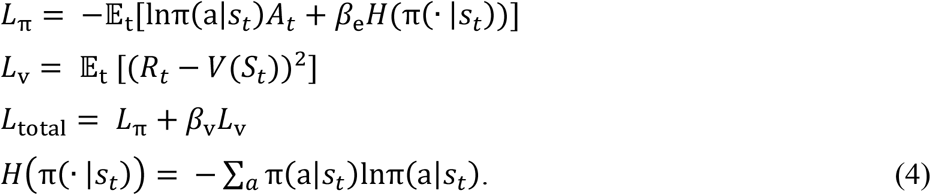

The gradients of *L*_total_ were backpropagated through time across the LSTMs.

A predefined outer-loop algorithm (first RL) was used to optimize network parameters, while training induced a distinct, learned RL procedure in the recurrent dynamics (second RL).^19^ The first RL was only for the training phase and did not determine choices during the test phase.

#### Hybrid deep reinforcement learning model

We designed a multiple-strategy agent of H-DRL based on the meta-RL framework, including the first RL (defined as weight RL) and the second RL (recurrent RL), with three key modifications:

1. Sequential online learning: weights were updated after each trial instead of being accumulated over multiple trials.
2. Positive activations: both activation functions and their derivatives remained positive.
3. Simple stochastic gradient descent (SGD) optimizer: SGD did not involve gradient accumulation or momentum.

The mathematical rationale of H-DRL is provided in the **Supplementary Note**. The RNN consisted of 100 fully connected units. Its dynamics were as follows:

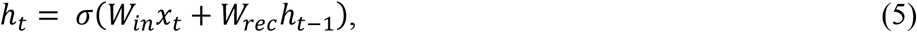

where *W*_*in*_ ∈ ℝ^D×100^ is the input weight matrix, *W*_*rec*_∈ ℝ^100×100^ is the recurrent weight matrix, *h*_*t*_ ∈ ℝ^100^ is the hidden activation at time step *t*, and *σ* denotes the activation function. We used the softplus activation function as *σ*.

The recurrent activities passed through a 20-unit fully connected layer and were mapped to two Q values (the expected reward of the selected action, given the current state):

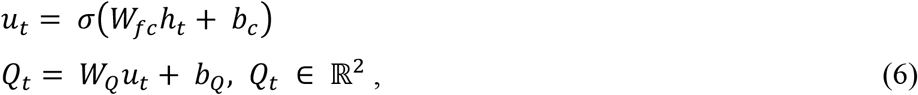

where *W*_*fc*_ ∈ ℝ^20×100^, *b*_*fc*_ ∈ ℝ^20^, *W*_*Q*_ ∈ ℝ^2×20^, and *b*_*Q*_ ∈ ℝ^2^. Actions *a*_*t*_ ∈ {0, 1} were sampled from the Boltzmann distribution over Q values:

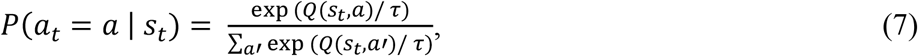

where temperature *τ* = 0.2. We used a simple per-trial Q-learning rule to improve interpretability and stability of learning compared to those of an actor-critic rule. At the end of trial *t*, the agent received the total reward. The instantaneous loss was

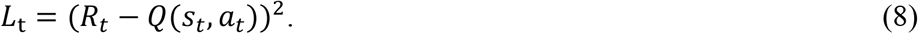

Immediately after each trial, parameters *θ* were updated via plain SGD and TBPTT:

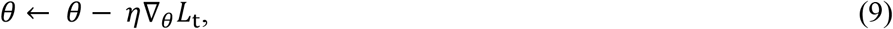

where *η* is the learning rate.

### Model simulation

We simulated both the meta-RL and H-DRL models on the trial structure used in our previous study in a perceptual decision-making task for a head-fixed mouse (**Figs. 2-5**).^15^ We also simulated choice behavior in a two-step decision-making task (**Fig. 1**).^3,10,11^

#### Perceptual decision-making task for head-fixed mice^15^

Before explaining the model simulation of our perceptual decision-making task, we briefly explain the task for mice. We used data from 26 male CBA/J mice (8–15 weeks old). We first trained mice to discriminate 90% low- (5–10 kHz) vs. high- (20–40 kHz) frequency tone clouds^31,33,34^ until the mice reached at least an 80% rate of correct choices. After the initial training, the mice either underwent a session in the neutral condition (transition probability (*p*) of 0.5) or proceeded directly to one of two probabilistic conditions of repeating (*p* = 0.2) or alternating (*p* = 0.9). The transition probability (*p*) defined the probability of switching the tone category of low or high in every trial. The repeating or alternating condition involved often repeating or switching tone categories, respectively.

Each trial began with retracting spouts away from the mice. After a random interval in the range of 1–2 s, the task presented a 0.6 s tone cloud. The mice then chose left or right spout after a delay of 0.5 s. Correct choices yielded 2.4 µL of 10% sucrose water, while incorrect choices triggered a 0.2 s noise burst. After at least 9 sessions in the repeating or alternating condition, the mice were introduced to an overtrained phase, where *p* = 0.2 and *p* = 0.8 were applied to the repeating and alternating conditions, respectively.

The overtrained phase included neural activity recording. Fourteen out of 26 mice were implanted with Neuropixels 1.0 probes targeting the orbitofrontal cortex (OFC), the striatum (STR), the motor cortex (M1), the posterior parietal cortex (PPC), the hippocampus (HPC), and the auditory cortex (AC).^15^ For the behavioral analyses, we used 51 and 127 sessions in the repeating and alternating conditions, respectively (**Fig. 2**). As in our previous study,^15^ we used the sessions for analyses when (i) the percentages of correct responses for both the 100% low tones and 100% high tones were greater than 75%, and (ii) the total reward amount in one session was at least 600 μL.

Among the recorded brain regions, we analyzed the neural data of OFC, PPC, HPC, and AC, where the data were obtained in both the repeating and alternating conditions (**Fig. 5**). For the neural data analyses, we added a criterion for the sessions in analyses in which the activity was simultaneously recorded from at least 50 neurons (repeating condition: OFC: 14 out of 17 sessions; PPC: 11 out of 18; HPC: 9 out of 18; AC: 15 out of 16; alternating condition; OFC: 18 out of 30; PPC: 16 out of 37; HPC: 17 out of 37; AC: 27 out of 31).

#### Simulation of the perceptual decision-making task

For the simulation of the models, each session consisted of 500 trials. Recurrent unit activities were initialized to zero at every beginning of each session. For simplicity, each trial was discretized into three steps: (1) ITI, (2) stimulus, and (3) decision (**Fig. 2a**).

The input vector *x*_*t*_ included the current stimulus (one dimension), the one-hot encoding of the previous action (0 or 1 with two dimensions), and the previous outcome (0 or 1 with one dimension; there were four input dimensions in total). The stimulus input was provided only at the stimulus step. Previous choices and outcomes were provided only at the ITI step. At the decision step, the output of the RNN was interpreted as a categorical choice (left or right). A reward was delivered immediately after the decision step with a magnitude of 1 for correct choices and 0 otherwise. The network maintained relevant information internally for subsequent steps.^21^

The stimulus input was a scalar representing the continuous-valued proportion of high-frequency tones, ranging between 0 and 1. In each trial, the mean proportion of the stimulus was determined based on the sound setting in our previous task,^15^ and the input was drawn from a truncated Gaussian distribution (SD = 0.25). For the neutral condition, the mean proportion of the stimulus was randomly selected from [0, 0, 0.2, 0.35, 0.65, 0.8, 1, 1]. For the repeating and alternating conditions, the mean proportion was probabilistically sampled from [0, 0, 0.25, 0.45] and [0.55, 0.75, 1, 1] for the low- and high-tone categories.

Additionally, we constructed an extended H-DRL model with eight steps per trial to analyze how the model memorized previous events during ITI (**Fig. 5b**). The extended model had five ITI time steps, one stimulus step, one delay step, and one decision step. The previous action and reward were provided to the model only at the first ITI step. The stimulus and decision steps were the same as in the original H-DRL model. The hyperparameters were identical to those of the original H-DRL model.

#### Model training and testing

Both meta-RL and H-DRL were first trained on the simulated task under the neutral condition (*p* = 0.5). We used the neutral condition until the rate of the model’s correct choices in one session reached 80 %. We then trained the model in either repeating or alternating conditions (*p* = 0.2 or 0.9) by using the pretrained weights on the neutral condition. Each model was trained for 20 sessions, followed by five overtrained sessions, in which the transition probability of the alternating condition was set to *p* = 0.8. This change in transition probability was the same as that in the mouse behavioral task.^15^ Ten independent instances of each model were trained. Each instance was initialized with different random weights to capture the variability in training dynamics.

For model testing, we did not freeze synaptic weights for either meta-RL or H-DRL even during the overtrained sessions. In the neutral condition, we analyzed the model choices during the last training session. In the repeating and alternating conditions, we analyzed all five sessions in the overtrained phase.

The hyperparameters of meta-RL were as follows: *lr* = 0.0005, γ = 0.5, *β*_e_ = 0.05, *β*_v_ = 0.05, and unroll length = 30 steps (10 trials). The hyperparameters of H-DRL were as follows: *lr* = 0.05, and unroll length = 3 steps (one trial). L2 regularization was used for H-DRL for continual learning.^66^ We set the decay rate to 0.001.

#### Two-step decision-making task

We validated the performance of H-DRL using a two-step task (**Fig. 1d**).^3,10,11^ The task design for model simulation was consistent with a previous study using meta-RL.^20^ In brief, at the first stage (*s*_0_), the model selected actions from two candidates (*a*_0_ or *a*_1_). Subsequently, the model transitioned to the second stage (*s*_1_ or *s*_2_) based on fixed transition probabilities. The transition structure was as follows:

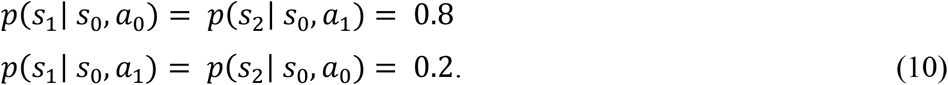

At the second stage, the model received an outcome based on the state and reward probability *P*_*rewarded*_:

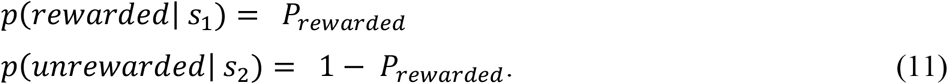

*P*_*rewarded*_ was either 0.1 or 0.9 and switched at the end of each trial with a probability of 0.025. The outcome was 1 (rewarded) or 0 (unrewarded). We trained 20 instances of the model independently. Each instance performed 100 sessions, which consisted of 100 trials per session. Next, we analyzed the choices in the last 25 sessions to validate model performance in the overtrained phase. The hyperparameters of H-DRL were the same as described above, except for *lr* = 0.1 and the decay rate of 0.01.

### Analysis of model-generated choices

#### Psychometric function

This section explains the analyses of discrete choices generated from the models in the perceptual decision-making task. The same analyses were already performed for the choices of the mice in our previous study.^15,30^

We first used a psychometric function in our previous study to analyze the generated choices.^30^ The psychometric function used a truncated Gaussian distribution between 0 and 1 for modeling sensory perception. The function had free parameters of stimulus sensitivity, choice bias, and lapse rates, which depended on the tone category in the previous trial. We used this function for the following analyses:

We used the full-parameter psychometric function to investigate whether the choices were biased by the previously rewarded choice in the neutral condition (**Fig. 2c**).

1. We used the full-parameter psychometric function to examine whether the choices were biased by the previous tone category in repeating and alternating conditions (**Fig. 2d left**).
2. We used a likelihood ratio test (p < 0.01) to identify the combinations of free parameters that explained the choices and examined when the previous-trial-dependent choice bias appeared (**Fig. 2d right**).
3. We also explored how the five most recent trials being correct or incorrect independently affected the choice in the current trial, together with the current sound stimulus (**Fig. 2e**). We used a logistic regression; the respective equation was reported in our previous study.^15,37^

#### Model-free and inference-based strategies

We investigated whether the model-generated choices matched a model-free or an inference-based strategy. We analyzed the choices with conventional RL models in our previous study^15^ (**Fig. 2f**). Both strategies hypothesized that mice estimated the hidden state of each trial (i.e., left-rewarded state (*S* = *L*) or right-rewarded state (*S* = *R*)) based on sensory stimuli. The model-free and inference-based strategies estimated the expected outcome and the state probability, respectively.

The model-free strategy used forgetting Q-learning^67^ and updated the expected outcomes of left and right choices in each state, defined as the prior value,^1,28,60^ while the belief state probability was fixed. The strategy used the prior values to compute a decision threshold in each trial with a softmax function. The decision threshold was then used to estimate the choice probability from the perceived sound modeled with a truncated Gaussian distribution. In contrast, when we applied such a model-free strategy to the RNN model, we removed the forgetting term, as previous research noted that forgetting Q-learning was inapplicable to deep-RL outputs due to outcome-independent choice biases.^21^

The inference-based strategy involved the belief state probability *P*(*S*) in each trial by estimating and updating the transition of state *P*_*transition*_ in every trial. The prior values were fixed. The decision threshold was computed with Bayesian inference based on *P*(*S*), and the choice probability was estimated the same way as for the model-free strategy. We updated *P*_*transition*_ based on the true state of the current and previous trials.

### Perturbation of H-DRL

We performed two perturbation tests to distinguish the contributions of weight- and recurrent-RLs in H-DRL in our perceptual decision-making task (**Fig. 3**):

- Weight-freeze test: All weight updates were frozen during test sessions.
- Activity-reset test: Between the ITI and stimulus steps in each trial, we reset all RNN units’ activities to zero, removing any across-trials memory in the recurrent activity.

### Analysis of recurrent dynamics of H-DRL

We explored how the internal representations of trained RNNs differed across task conditions. To this end, we performed the following encoding and decoding analyses of units’ activity (**Fig. 4**). In these analyses, we focused on four different time points of RNNs (**Fig. 4a**): (1) the initial random network, (2) after being trained with the neutral condition, (3) after being trained with the repeating condition, and (4) after being trained with the alternating condition. At each time point, we froze all weight updates, set the transition probability to 0.5, and performed five testing sessions to investigate the intrinsic network dynamics.

#### Encoding analysis of model activity

A generalized linear model (GLM) was used to analyze whether the RNN unit activity at the decision step represented the current choice (*C*), the current stimulus (*S*), the previous choice, the previous stimulus, and the previous outcome (*O*) (**Fig. 4b**) (Python, statsmodels.api. GLM, Gaussian family):

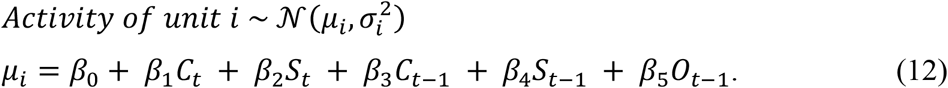

We first fit each unit activity to the full GLM using 10-fold cross validation (CV) repeated 100 times. To identify the task variables that were significantly represented in each unit, we applied a drop-one analysis.^15^ A reduced model was generated by omitting a target variable, and the same CV procedure with 100 repetitions was performed. If the average deviation of the full model was significantly lower than the mean minus 1.96 standard deviations of the omitted-variable model, the omitted target variable was defined as represented in the unit. This procedure was repeated for all variables and all units.

#### Decoding analysis of model activity

From the RNN unit activities at the decision step, we decoded the binary category of (1) the current stimulus *S*_*t*_ or (2) the previous stimulus *S*_*t*−1_ with a linear support vector machine (**Fig. 4c**).^38^ To prevent crosstalk of decoding performance between the previous and current stimuli, we randomly subsampled the trials so that the numbers of trials for the four combinations of previous and current stimuli were equal. To account for randomness due to subsampling and fold assignment, we repeated subsampling and 10-fold nested CV 100 times and used the average rate of correct choices as the decoding performance. We also investigated the absolute cosine similarity between the decoding vectors of *S*_*t*_ and *S*_*t*−1_.

#### Analysis of gradient-retention dynamics in RNNs

To quantify whether RNNs with identical initial weights exhibit rich or lazy learning depending on task conditions, we visualized how the gradient-retention capacity evolved over the course of training (**Fig. 4f**). While linear RNNs can be evaluated using the spectral radius as a global measure of memory retention, nonlinear RNNs require a more localized approach based on the Jacobian matrix at specific input states.^39^

Given that our networks were trained using BPTT for three steps, we computed the chain Jacobian over three consecutive time steps within each episode. For each hidden state *h*, we computed the derivative of the softplus activation, which corresponded to the sigmoid function *σ*(*z*) = 1/(1 + *e*^−*z*^), and constructed the diagonal matrix *D* = *diag*(*σ*(*z*)). The local Jacobian was given by

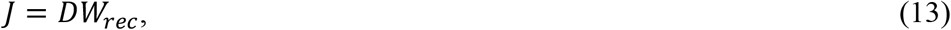

where *W*_*rec*_ is the recurrent weight matrix. Because we updated the weights after every trial (three steps), we obtained the chain Jacobian by multiplying the local Jacobians over three time steps:

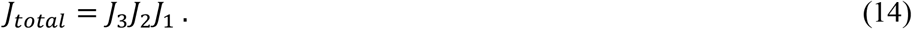

Finally, we computed the spectral radius of the chain Jacobian, defined as the maximum absolute eigenvalue:

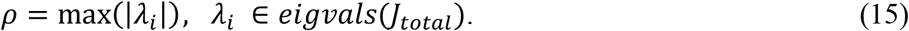

This spectral radius served as an index of the network’s ability to retain input information.

### Comparison of activity between mouse neurons and RNN units

#### Decoding of task variables during ITI

We decoded task variables from the activity of mouse neurons and that of model units during the inter-trial interval using L1-regularized sparse logistic regression (**Fig. 5c, d**). The decoded variables were the previous choice, the correct side, and the outcome. To prevent crosstalk of decoding performance among the three variables, we randomly subsampled the trials so that the numbers of trials for the four combinations of previous choices and outcomes were identical.

For the unit activity of H-DRL and meta-RL, all 100 units were included. For the neural data, we analyzed the data from OFC, PPC, HPC, and AC in which neural activity was recorded in both the repeating and alternating conditions. In each decoding analysis, 50 neurons were randomly subsampled. We repeated subsampling and 10-fold nested CV 100 times and used the average rate of correct choices as the decoding performance.

#### Projection of activity to the choice axis

To explore the population activity of OFC, we projected the activity during −2.0 – 1.1 s relative to the onset of sound to a choice axis (**Fig. 5e**). The choice axis was defined using the activity during the delay period (0.6–1.1 s from the onset of sound). We used the SLR with 10-fold CV for the choice decoder, and the weight vector of the SLR was used as the projection axis. We visualized four conditions of neural activity defined by the combinations of correct previous and current sides. To prevent the effects of a biased number of trials depending on the previous outcome (0 or 1), the projection axis of the previous correct side was computed separately for each previous outcome and averaged across the two cases.

## Quantification and statistical analysis

We used Python 3.11.4 and MATLAB 2022b for all analyses. The solid lines and shaded areas in the figures represent the means ± standard deviations (SD) or standard errors (SEM). Model fitting with conventional RL was performed with the Bayesian information criterion (BIC) (**Fig. 2f**). For the behavioral analyses of multiple sessions of each mouse, we used a linear mixed-effects model (MATLAB: fitlme).^15^ We mainly used two-sided nonparametric statistical tests. Descriptions were made for one-sided statistical tests. All statistical details are provided in figure legends, figures themselves, or the descriptions of the results.

## Data availability

All simulation data generated in this study will be made available upon publication.

The behavioral and electrophysiological data from our previous study are available at https://doi.org/10.17632/vf4b4bmzjp.1.

## Code availability

The code is available at https://github.com/hayato0405/Maeda_et_al.

## Acknowledgments

This research was funded by JSPS Kakenhi (Grant Nos. JP24H02150, JP24KK0186, JP21H05243, JP23K21693, JP25H02599, JP25H01130), AMED (Grant Nos. JP24wm0625415, JP24gm6510019), the Uehara Memorial Foundation, the Narishige Neuroscience Research Foundation, and the Takeda Science Foundation for A.F.

## Author contributions

H.M. analyzed the data and proposed the model. S.W. collected the animal data. H.M. and A.F. designed the study and wrote the paper.

