## Supplementary Note and Figures for "Dual reinforcement-learning network modules for modeling decision-making with multiple strategies"

#### 1. Objectives

This **Supplementary Note** provides the mathematical and theoretical details that complement the main text. The method proposed in this study, called "hybrid deep reinforcement learning" (H-DRL), extends the canonical $RL^2$-based meta-reinforcement learning (Meta-RL) framework with minimal modifications to serve as a theoretical model of multiple-strategy implementation.

**Section 2** presents the background, including the mathematical formulation of $RL^2$-based Meta-RL and the reasons it is insufficient to account for multiple learning strategies. **Section 3** introduces the core mechanisms of H-DRL and explains the rationale for their design. Finally, **Section 4** details the architectural components implemented in our experiments, clarifying the design considerations underlying each choice.

#### 2. Background

##### 2-1. Overview

In this section, we outline the theoretical background that underpins our proposed framework and discuss the limitations of existing approaches. Our theory builds upon the $RL^2$ architecture, a meta-reinforcement learning model implemented with deep recurrent networks. $RL^2$ has recently gained attention as a computational model of the prefrontal and orbitofrontal cortices,[1–3] and its strengths as a cognitive model have been extensively discussed.[4] However, comparatively little attention has been devoted to its limitations, both as a learning algorithm and as a model of biological cognition.

In **Section 2-2**, we first present the mathematical formulation of the $RL^2$-based Meta-RL framework. Next, **Section 2-3** reviews previous theoretical research interpreting $RL^2$ as an implicit approximation of Bayesian optimal inference. Finally, in **Section 2-4**, we examine the strengths and limitations of $RL^2$ used as a cognitive model, and highlight the theoretical gap that motivates our proposed extension.

##### 2-2. Mathematical Foundation of $RL^2$-based Meta-RL

We begin by formalizing the canonical $RL^2$ framework, which provides the mathematical foundation for most recurrent meta-reinforcement learning approaches.[5,6] The central idea of $RL^2$ is to enable rapid task adaptation through the evolution of an internal recurrent state rather than by re-optimizing policy parameters from scratch for each new environment. This allows an agent to utilize its interaction history directly to guide future behavior.

We consider a distribution of partially observable Markov decision processes (POMDP), defined as

$$T_\phi = (S, A, O, P_\phi, \Omega_\phi, R_\phi, \gamma) \tag{1}$$

where $\phi \sim p(\phi)$ indexes a specific task drawn from a task distribution $p(\phi)$. Above, $S$ denotes the set of latent states, $A$ is the action space, and $O$ represents the observation space.

The environmental dynamics are described by the transition kernel $P_\phi(s_{t+1}|s_t, a_t)$, the observation model by $\Omega_\phi(o_t|s_t)$, and the reward function by $R_\phi(s_t, a_t)$. The discount factor is denoted by $\gamma \in (0, 1)$.

At each time step $t$, the agent receives an observation $o_t \in O$, selects an action $a_t \in A$, transitions to a new state $s_{t+1}$, and receives a scalar reward $r_t \in R$. Since the task parameter $\phi$ is unknown to the agent, the latter must adapt its decision-making strategy based solely on the sequence of past interactions.

In the RL² framework, the policy is often parameterized as a recurrent neural network (RNN) that maintains a hidden state $h_t \in \mathbb{R}^d$ as a compact representation of the agent's interaction history. The hidden state evolves according to

$$h_{t+1} = f_\theta(h_t, x_{t+1}) \tag{2}$$

where $f_\theta$ is a recurrent update function (e.g., GRU or LSTM) parameterized by $\theta$. The input vector

$$x_{t+1} = [o_{t+1};\ a_t;\ r_t] \tag{3}$$

concatenates the new observation, the previous action, and the received reward. The policy itself is defined as a conditional distribution over actions given the hidden state:

$$\pi_\theta(a_t|h_t) = g_\theta(h_t) \tag{4}$$

where $g_\theta$ maps the hidden state to action probabilities.

During meta-training, the parameter $\theta$ is optimized across multiple tasks sampled from $p(\phi)$ to maximize the expected cumulative return:

$$\max_\theta \mathbb{E}_{\phi \sim p(\phi)} \mathbb{E}_{\tau \sim \pi_\theta, T_\phi} \left[\textstyle\sum_{t=0}^{T} \gamma^t r_t\right] \tag{5}$$

where $\tau = \{(o_t, a_t, r_t)\}_{t=0}^{T}$ denotes a trajectory collected under policy $\pi_\theta$. Importantly, once meta-training is complete, parameters $\theta$ remain fixed during deployment on new tasks. When a novel task is encountered, all adaptation is achieved through the online evolution of the hidden state $h_t$, which is continuously updated as the agent interacts with the environment.

##### 2-3. Meta-RL as a Bayesian Inference Approximator

Prior theoretical research[7] frames meta-RL (e.g., RL²) as an **amortized Bayesian posterior prediction** over a task distribution. During deployment, the network parameters $\theta$ are fixed; adaptation proceeds via updates of the recurrent state $h_t$ driven by past interactions.

Consider a mixture over interaction sequences $ao_{\le T} = (o_1, a_1, \dots o_T, a_T)$:

$$P(ao_{\le T}) = \textstyle\sum_\phi P(ao_{\le T}|\phi)p(\phi) \tag{6}$$

This is a marginal data-generating distribution rather than a policy definition; it serves as a basis for posterior prediction.

For action selection, Thompson sampling draws from the posterior predictor:

$$a_{t+1} \sim P(a_{t+1}|\hat{a}o_{\leq t}) = \sum_{\phi} P(a_{t+1}|\phi, ao_{\leq t})P(\phi|\hat{a}o_{\leq t}) \quad (7)$$

Here, the "hat" notation indicates a causal intervention on past actions. The posterior is updated recursively as

$$P(\phi|\hat{a}o_{\leq t}) = \frac{P(o_t|\phi, ao_{<t})P(\phi|\hat{a}o_{<t})}{\sum_{\phi'} P(o_t|\phi', ao_{<t})P(\phi'|\hat{a}o_{<t})} \quad (8)$$

To amortize the computation of the posterior predictor in (7)–(8), the recurrent update and policy readout defined in **Section 2-2** are used, where the hidden state $h_t$ serves as a compact summary of past interactions, and $g_\theta(h_t)$ approximates the posterior predictive distribution in (7).
From the sequential-prediction perspective, Thompson sampling is compression-optimal under the logarithmic loss, and meta-training minimizes the expected logarithmic loss. In practice, we use the Monte-Carlo approximation

$$\mathbb{E}[\ell] \approx \frac{1}{N}\sum_{n=1}^{N} \ell(f_\theta; \tau^{(n)}) \quad (9)$$

with trajectories $\{\tau^{(n)}\}$ sampled from the task prior $p(\phi)$ during meta-rollouts; at deployment, $\theta$ remains fixed, and adaptation occurs through $h_t$.

#### 2-4. Gap between animals and Meta-RL agents

The mathematical formulation above highlights an important implication: within the canonical $RL^2$ framework, the agent continually updates its internal state $h_t$ through an activity-based reinforcement learning process. This mechanism, which incrementally integrates observations, actions, and rewards, can in theory converge toward a near-Bayesian optimal strategy over latent task variables. From a computational neuroscience perspective, such activity-driven adaptation corresponds to the **model-based** or **inference-based** strategy—generalized to continuous state and observation spaces. Unlike classic model-based approaches that presuppose a fixed environmental model, $RL^2$ implicitly constructs and updates an internal model through experience, thereby providing a powerful unifying account of model-based behavior.

However, a fundamental gap remains between Meta-RL agents and animal behavior. Empirical and theoretical studies have long emphasized that animals flexibly deploy and integrate multiple learning systems—both goal-directed (model-based) and habitual (model-free) strategies—depending on task demands.[8–12] The $RL^2$ architecture, while capable of representing inference-based adaptation, relies on a single recurrent update mechanism and thus cannot explicitly capture the coexistence or arbitration between qualitatively distinct learning modes. In other words, its adaptation dynamics are monolithic: all forms of flexibility must emerge from the same recurrent pathway, without an explicit mechanism for switching or blending multiple strategies.

Beyond this theoretical limitation, there are also engineering constraints. First, $RL^2$ agents are fragile to out-of-distribution (OOD) shifts. When such agents face tasks that deviate from the training distribution $p(\phi)$, the internal dynamics learned during meta-training fail to extrapolate, often collapsing toward suboptimal or random behavior. Second, the meta-learning process itself is computationally slow and sample-inefficient since each meta-update requires averaging over all reinforcement learning episodes nested within the outer optimization loop. These issues are typically masked if performance is evaluated only by post-training test accuracy, but become critical in dynamic environments where animals must maximize reward from the very beginning of interaction, without extensive pretraining.

Overall, these observations suggest that while $RL^2$ provides an elegant formalization of inference-based adaptation, its structure remains too rigid to reproduce the dual-system flexibility observed in biological agents.

#### 3. Proposed Framework

##### 3-1. Model Overview

As discussed in **Section 2**, $RL^2$-based Meta-RL can autonomously acquire model-based behavior and has been proposed as a biologically plausible model of prefrontal and orbitofrontal function. Building on this framework, our goal is to reproduce the flexible integration and switching of multiple strategies observed in animals while maintaining minimal structural complexity. The latter is essential for interpretability and biological plausibility.

While Meta-RL naturally develops model-based behavior, it lacks an explicit mechanism for model-free control. One straightforward approach would be to append an external model-free reinforcement learning (RL) module.[13,14] However, this ad hoc addition would obscure how the two systems arbitrate and would compromise the parsimony of the model.

Instead, we propose a more intrinsic and principled extension called "hybrid deep reinforcement learning" (H-DRL). Our key insight is that $RL^2$ already embeds a model-free component within its architecture. In $RL^2$, a conventional RL rule defines the outer-loop loss, which trains the recurrent network to establish a distinct, autonomous inner-loop RL process. Thus, classic RL is not absent from $RL^2$; it merely remains latent, confined to shaping learning signals rather than behavior itself.

In standard implementations, model parameters are updated infrequently (e.g., per session), rendering this model-free pathway behaviorally silent. We hypothesize that if such weight updates are performed online, on a trial-by-trial basis, the same gradient updates that once served as learning signals will directly influence behavioral policy. In this way, $RL^2$ can express a model-free learning channel without any architectural addition, simply by reactivating an element already inherent in its design.

Accordingly, H-DRL introduces only three minimal modifications to the canonical $RL^2$ framework, each preserving its structure while unlocking the latent dual-learning mechanism described above.

**1. Trial-by-trial weight update.**

Network parameters are updated immediately after each trial rather than after full task episodes.

This modification converts the outer-loop optimization of $RL^2$ into an *online reinforcement process* operating at the behavioral timescale, allowing weight changes to directly influence subsequent actions. Consequently, the weight space itself functions as a model-free reinforcement channel (**weight-RL**) that coexists with the recurrent, inference-based adaptation (**recurrent-RL**).

**2. Positive activations in the recurrent network**

All recurrent units are constrained to strictly positive activations,

$$\forall i, t:\ h_{t,i} > 0 \tag{10}$$

This constraint prevents biologically unrealistic behavior inherent in standard zero-centered activation functions (e.g., tanh). In such standard models, a single unit can output both positive and negative values, effectively switching between excitatory and inhibitory roles depending on its input. However, biological neurons are strictly distinct in their effect (Dale's principle). By enforcing sign-restricted activity, our model ensures that the functional role of each unit remains consistent, aligning with these fundamental neuronal properties.

**3. Pure SGD**

Parameter updates follow a vanilla stochastic gradient descent (SGD) rule,

$$\theta_{t+1} = \theta_t - \eta \nabla_\theta L_t \tag{11}$$

with no momentum, gradient accumulation, or adaptive rescaling (e.g., Adam or RMSProp). Each update thus depends solely on the current reward-prediction error, which is consistent with the dopamine-driven, trial-by-trial synaptic plasticity observed in biological reinforcement learning.

These modifications are minimal yet sufficient to expose the dormant dual-learning structure of $RL^2$. As we demonstrate in the following sections, they endow the model with three key properties: (1) coexistence of weight-RL and recurrent-RL within a single network, (2) the emergence of model-free behavioral dynamics in weight-RL, and (3) spontaneous strategy switching between the two systems without any explicit arbitrator.

##### 3-2. Formalization of Concurrent Adaptation

To formalize the coexistence of two adaptation processes—recurrent-RL through $h_t$ and weight-RL through $\theta_t$—we extend the canonical $RL^2$ framework by allowing both to evolve online within a single architecture. The goal of this derivation is to express, in a unified mathematical form, how recurrent-state dynamics and synaptic weight changes jointly affect the policy output.

At each time step $t$, corresponding to one behavioral cycle (observation $o_t$, action $a_t$, and reward $r_t$), parameters are updated as follows:

$$\theta_{t+1} = U(\theta_t, h_t, a_t, r_t) \tag{12}$$

where $U$ is a **fixed** update operator (e.g., a predefined RL rule). The hidden state subsequently evolves under the new parameters,

$$h_{t+1} = f_{\theta_{t+1}}(h_t, x_{t+1}) \tag{13}$$

The policy output at each step is

$$u_t = \pi_{\theta_t}(h_t) = g_{\theta_t}(h_t),\ u_{t+1} = \pi_{\theta_{t+1}}(h_{t+1}) = g_{\theta_{t+1}}(h_{t+1}) \tag{14}$$

We define the first-order change in the policy output as

$$\Delta u_t \coloneqq u_{t+1} - u_t \tag{15}$$

To isolate how both adaptation channels contribute to this change, we perform a first-order Taylor expansion around the current variables $(h_t, \theta_t)$.

Expanding $h_{t+1}$ with respect to the weight update $\Delta\theta_t \coloneqq \theta_{t+1} - \theta_t$ yields

$$h_{t+1} = f_{\theta_t}(h_t, x_{t+1}) + J_\theta^f \Delta\theta_t + O(\|\Delta\theta_t\|^2),$$
$$J_\theta^f \coloneqq \frac{\partial f_\theta(h_t, x_{t+1})}{\partial\theta} \Big|_{\theta_t} \tag{16}$$

This step separates the contribution of the recurrent update from that of the weight change. Substituting (16) into the policy output and applying the chain rule leads to

$$\Delta u_t \approx J_h^\pi\big(f_{\theta_t}(h_t, x_{t+1}) - h_t\big) + (J_h^\pi J_\theta^f + J_\theta^\pi)\Delta\theta_t + R_t \tag{17}$$

where

$$J_h^\pi \coloneqq \frac{\partial \pi_\theta(h)}{\partial h} \Big|_{(h_t,\theta_t)},\ J_\theta^\pi \coloneqq \frac{\partial \pi_\theta(h_t)}{\partial\theta} \Big|_{\theta_t} \tag{18}$$

Each Jacobian quantifies the local sensitivity of the policy to hidden-state and parameter changes.

The residual term represents a collection of higher-order effects,

$$R_t = O\begin{pmatrix} \left\|f_{\theta_t}(h_t, x_{t+1}) - h_t\right\|^2 + \|\Delta\theta_t\|^2 + \\ \|\Delta\theta_t\|\left\|f_{\theta_t}(h_t, x_{t+1}) - h_t\right\| \end{pmatrix} \tag{19}$$

This formulation expresses the first-order policy variation as a linear superposition of activity-driven and weight-driven components, clarifying that within the local approximation, the two adaptation processes coexist as separable yet concurrent contributions.

##### 3-3. Mechanistic Basis of "Model-free behavior"

###### 3-3-1. "Model-free behavior" in neuroscience

In reinforcement learning theory, the term "model-free" refers to algorithms that update action values directly from reward-prediction errors without using an explicit model of environmental transitions. In neuroscience, however, the term is used **operationally** to describe a specific

behavioral signature: the reinforcement or the weakening of state–action pairs based on outcome valence (the Law of Effect), often manifesting as habitual or stimulus–response control.
This distinction becomes evident in the two-step task, where behavioral analyses classify control strategies as "model-free" or "model-based" based on the pattern of action reinforcement (specifically, the presence or absence of a reward × transition interaction). However, this behavioral taxonomy—derived from canonical algorithms such as SARSA and REINFORCE—does not align perfectly with the algorithmic definition based on model usage. Meta-RL clearly illustrates this divergence: although architecturally model-free, it exhibits fully model-based behavior in the two-step task.[1]
In what follows, we therefore use the term "model-free behavior" in the operational sense, i.e., defined by monotonic updates of action values according to the sign of the instantaneous reward-prediction error (RPE), independent of any explicit environmental model.

##### 3-3-2. Formal derivation of model-free constraints in weight-RL

To formally demonstrate that the weight-based update channel (weight-RL) yields model-free behavior in the operational sense defined above, we analyze the gradient structure under the architectural constraints introduced in **Section 3-1**.
Here, we focus our analysis on the updates applied to the linear readout that maps hidden activity $h_t$ to the policy output:

$$u_t = W^{(L)}h_t + b^{(L)} \tag{20}$$

Given the scalar loss $L(u_t)$ defined by RPE, the gradients are

$$\frac{\partial L}{\partial W_j^{(L)}} = \frac{\partial L}{\partial u_t}h_{t,j},\ \frac{\partial L}{\partial b^{(L)}} = \frac{\partial L}{\partial u_t} \tag{21}$$

Since $h_{t,j} > 0$, the sign of every gradient component is determined solely by the sign of the output error:

$$sgn\left(\frac{\partial L}{\partial W_j^{(L)}}\right) = sgn\left(\frac{\partial L}{\partial b^{(L)}}\right) = sgn\left(\frac{\partial L}{\partial u_t}\right) \tag{22}$$

Accordingly, a one-step SGD update with a learning rate $\eta > 0$,

$$\Delta W_j^{(L)} = -\eta\frac{\partial L}{\partial u_t}h_{t,j},\quad \Delta b^{(L)} = -\eta\frac{\partial L}{\partial u_t} \tag{23}$$

applied to the output weights, moves all parameters in the readout layer uniformly in the same direction as the sign of RPE. A positive RPE increases subsequent outputs across all inputs, whereas a negative RPE uniformly reduces them. Under positive activations and memoryless SGD, this produces a monotonic, sign-consistent modification of the readout weights directly driven by instantaneous RPEs.
Theoretically, such strict component-wise sign preservation is guaranteed at the linear readout. However, empirically, if smooth nonnegative activations are used, we observe that "value inversion"—where a positive RPE leads to a decrease in Q-values—does not occur even if updates are backpropagated through nonlinear layers. This makes the system functionally equivalent to a globally monotonic, RPE-driven process that instantiates the computational hallmark of model-free reinforcement defined in **Section 3-3-1**.

#### 3-4. Natural arbitration between weight-RL and recurrent-RL

H-DRL integrates two concurrent learning channels, namely, weight-RL, producing model-free behavior, and recurrent-RL, implementing inference-based adaptation.
Beyond their coexistence, the model exhibits natural arbitration between the two, switching dominance depending on task statistics.
Unlike classic dual-strategy models with an explicit mixing parameter, here arbitration arises from the asymmetry between the two processes. Weight-RL operates from the onset of learning but lacks flexibility, whereas recurrent-RL, driven by meta-learning of the recurrent network, evolves gradually as experience accumulates. Thus, early behavior is dominated by weight-RL, while extended training can shift control toward recurrent-RL.
Importantly, because H-DRL performs online updates, it directly receives the temporal structure of task statistics as a bias in gradient noise; this is a structure that would otherwise be lost in batch optimization. We hypothesize that temporal correlations in this noise modulate the effective learning rate and thereby alter the choice of the dominant reinforcement process. If consecutive updates are positively correlated, weight-RL gains a transient advantage; if they are anti-correlated, meta-learning of recurrent-RL is facilitated. This mechanism yields spontaneous, environment-dependent switching between the two strategies without any external arbitration.

We formalize this intuition in a simplified setting. Specifically, we analyze how the temporal correlation structure of gradient noise affects the asymptotic behavior of online parameter updates. The goal is to characterize how these correlations determine the effective learning rate, the steady-state error, and the speed of convergence without invoking any explicit arbitration or control mechanism.
Let $e_t = \theta_t - \theta^*$. Near the optimum, the linearized recursion is

$$e_{t+1} = (I - \eta_t H)e_t - \eta_t \zeta_t \tag{24}$$

where $H = \nabla^2 F(\theta^*) \succ 0$ with eigenvalues $\mu \leq \lambda_i \leq L$.
The noise sequence $\{\zeta_t\}$ is assumed to be conditionally unbiased, given the parameters, $E[\zeta_t|\theta_t] = 0$, weakly stationary, and square-integrable. Its autocovariance and spectrum are

$$\Gamma(k) = E[\zeta_t \zeta_{t-k}^\top], \; S_\zeta(\omega) = \frac{1}{2\pi}\sum_k \Gamma(k)\, e^{-i\omega k} \tag{25}$$

The long-term covariance,

$$\Lambda = \sum_k \Gamma(k) = 2\pi S_\zeta(0) \tag{26}$$

is finite for short-memory noise and divergent for long-memory processes. Short-lag positive correlations increase $\Lambda$, while negative correlations reduce it.

##### Constant step size

For a constant $\eta_t \equiv \eta$ with $0 < \eta < 2/L$, define the stationary covariance $\sum_e = lim_{t \to \infty} E[e_t {e_t}^\top]$.
The leading-order solution of the discrete Lyapunov equation gives

$$\textstyle\sum_e = \eta L_H^{-1}(\Lambda) + O(\eta^2), \; L_H(X) = HX + XH \tag{27}$$

if $H$ and $\Lambda$ commute,

$$\textstyle\sum_e = \frac{\eta}{2} H^{-1}\Lambda + O(\eta^2) \tag{28}$$

The steady-state parameter mean-square error and function-value gap are therefore

$$E_\infty = tr(\Sigma_e) = \frac{\eta}{2} tr(H^{-1}\Lambda) + O(\eta^2)$$
$$F_\infty = \frac{1}{2} tr(H\Sigma_e) = \frac{\eta}{4} tr(\Lambda) + O(\eta^2) \quad (29)$$

In one dimension, these reduce to $E_\infty = \eta\Lambda/(2\lambda)$ and $F_\infty = \eta\Lambda/4$.

**Decaying step size**

For a decaying step size $\eta_t = c/t$ with $c > 1/(2\mu)$, let $H = Qdiag(\lambda_i)Q^\top$ and $\Lambda' = Q^\top \Lambda Q$, with diagonal elements $\Lambda_i = (\Lambda')_{ii}$.

Then, under short memory,

$$\lim_{t\to\infty} tE[\| e_t \|^2] = \sum_{i=1}^{d} \frac{c^2}{2c\lambda_i - 1} \Lambda_i$$
$$\lim_{t\to\infty} tE[F(\theta_t) - F(\theta^*)] = \frac{c^2}{2} \sum_{i=1}^{d} \frac{\lambda_i}{2c\lambda_i - 1} \Lambda_i \quad (30)$$

Thus, the convergence rate remains

$$E[F(\theta_t) - F(\theta^*)] = \frac{c^2}{2} \sum_{i=1}^{d} \frac{\lambda_i}{2c\lambda_i - 1} \Lambda_i \frac{1}{t}\big(1 + o(1)\big) = \Theta(1/t) \quad (31)$$

but the coefficient depends jointly on the curvature spectrum $\{\lambda_i\}$ and the noise correlation structure through $\Lambda_i$.

If noise instead exhibits long memory, $\Gamma(k) \sim Ck^{-\alpha}$ with $0 < \alpha < 1$, the order itself degrades:

$$E[F(\theta_t) - F(\theta^*)] = \Theta(t^{-\alpha}),\ E[\| e_t \|^2] = \Theta(t^{-\alpha}) \quad (32)$$

with the borderline case $\alpha = 1$ yielding $(log t)/t$.

**Interpretation**

Our analysis demonstrates that temporal correlations in gradient noise significantly modulate the effective learning dynamics across both noise regimes.

Under short-memory conditions, while the asymptotic convergence rate remains theoretically invariant, the correlations directly scale the pre-factor of parameter fluctuations. Thus, the steady-state variance is amplified or suppressed by the noise structure $\Lambda$, effectively altering the "temperature" of the optimization. Under long-memory conditions, the effect is even more pronounced, as correlations alter the convergence rate itself, producing sublinear adaptation.

In both cases, the physical implications are consistent: positive correlations maintain higher volatility in the weight-RL channel (slowing convergence or increasing variance), whereas negative correlations promote stability. Thus, regardless of the specific memory structure of noise, temporal correlations provide a robust mechanistic basis for the spontaneous arbitration between weight-RL and recurrent-RL.

##### 3-5. Summary

As shown above, H-DRL is a minimal extension of $RL^2$, yet it has strong theoretical utility as a model that explains multiple-strategy implementation. H-DRL offers an alternative to the classic dual-strategy model[8] in which a model-free learner and a model-based learner operate as independent computational modules and are integrated by an external arbitrator. Although this model has successfully explained animal behavior and has been widely accepted as a major theoretical framework, it faces two major issues: (i) experimental evidence does not fully support the hypothesis that model-free and model-based computations occur in distinct brain regions, and (ii) the neural basis of the external arbitrator remains elusive. Recent studies therefore suggest that model-free and model-based processes may be more integrated than previously thought.

H-DRL provides a new perspective: trial-by-trial weight updates correspond to model-free behavior, while the long-term accumulation of such weight changes, in conjunction with neural activity, corresponds to model-based behavior. Thus, two strategies naturally emerge within a single circuit, and integration or switching occurs through interaction with the task. This framework offers a new view that the brain may achieve flexible multi-strategy learning not by maintaining two independent systems but through a unified reward-based learning process.

### 4. Discussion of architectural choices

#### 4-1. Overview

In this section, we discuss the rationale of the architectural design of our model. Ideally, comparisons with standard $RL^2$ should involve modifications only to the core components described above. However, several practical adjustments were necessary due to implementation issues inherent in online learning. In addition, the overall architecture of H-DRL was determined from multiple perspectives, not limited to technical feasibility. The following subsections describe these considerations in detail.

#### 4-2. Training mechanism: BPTT or RTRL

As noted in the **Discussion** section of our main text, BPTT has several limitations. First, the backpropagation framework has long been criticized for its limited biological plausibility.[15] Second, its performance is highly sensitive to the choice of the context window.[16]

An alternative direction is to use real-time recurrent learning (RTRL),[17] which is inherently suitable for online updates and, in principle, capable of capturing dependencies that extend beyond the truncated windows of BPTT. Moreover, biologically motivated variants such as RFLO[18] have been proposed, and RFLO has recently been applied to deep reinforcement learning tasks.[19]

Despite these attractive properties, we opted to use BPTT in this study. The primary reason is that BPTT remains theoretically well understood and widely used in existing meta-RL research, facilitating direct comparisons with prior analyses. Importantly, earlier studies do not entirely dismiss the biological plausibility of backpropagation-like mechanisms.[15] In contrast, RTRL suffers from prohibitive computational costs, and its theoretical and empirical foundation is still comparatively underdeveloped. Although RTRL and its variants represent a promising avenue for future extensions of our model, we refrained from adopting them in the present study due to these unresolved challenges.

#### 4-3. Outer-loop learning rule: Actor-critic or Simple Q-learning

In $RL^2$-based meta-reinforcement learning, the outer loop is often optimized with actor–critic algorithms such as TRPO,[5] A2C,[2,6] or ACER.[20] In contrast, we used a simple Q-learning update

for the outer loop. The rationale is as follows.
First, recent studies have experimentally demonstrated that modern deep policy gradient methods tend to fail catastrophically under sequential online learning conditions.[21] It is well known that actor–critic methods become unstable when updated in a strictly one-step online process, and we also observed divergence in our own experiments when using such methods.
Second, actor–critic architectures are not well suited for continual learning. H-DRL is designed under the assumption that parameter updates continue throughout both the learning and overtrained phases. However, in an actor–critic architecture, the actor's output distribution $p(a_t|h_t)$ often collapses toward nearly deterministic values (close to 0 or 1) as training progresses, regardless of the critic's reward prediction error. As a result, the policy gradient almost vanishes, and further adaptation becomes ineffective.
Finally, reading both the actor and critic outputs from the same recurrent network complicates the interpretation of its internal representations, whereas separating the two networks would make it difficult to maintain interpretability from the perspective of a biological or cognitive model. For these reasons, we adopted a simple Q-learning formulation, which preserved stability and interpretability in an online and continual learning setting.

#### 4-4. Reward prediction error formulation

In classic reinforcement learning, the temporal difference (TD) error—typically formulated as TD(0) or TD(λ)—is a natural choice for defining the reward prediction error in online learning. However, in our setting, where each trial consists of multiple steps, as in standard deep reinforcement learning simulations, computing the value of the next trial at the time of weight updates poses a unique challenge.
In particular, under a partially observable Markov decision process, the next state $s_{t+1}$ is not explicitly available to the recurrent network; it can only be inferred through predictions based on the input distribution, which itself is unknown to the model. As a result, such estimation is practically infeasible.
Therefore, acknowledging this simplification, we adopted the following minimal form of reward prediction error:

$$\delta = r_t - Q(s_t, a_t) \tag{33}$$

Notably, methods such as true online TD(λ)[22] allow computation of TD errors following an online backward-view procedure without looking ahead into future rewards. This method ensures consistency between forward and backward views via eligibility traces and has been validated in the context of linear function approximation. However, its theoretical extension to nonlinear approximators remains limited, the implementation is complex, and the interpretability varies depending on the choice of λ. For these reasons, we prioritized simplicity and interpretability and did not use this approach in the present study.

#### 4-5. Activation function

Among the activation functions commonly used in deep learning, several produce strictly nonnegative outputs; such functions include sigmoid, softplus, and ReTanh. We adopted the softplus function, which has been frequently used in neuroscientifically grounded RNN models.[23,24] Investigating how the model behaves under alternative nonnegative activations such as sigmoid or ReTanh would be an interesting direction for future research.
Relaxing the sign constraint to allow zero-valued activations would make ReLU an attractive

option, given its widespread use. However, ReLU units suffer from complete gradient blockage once their activation becomes zero, effectively partitioning the network into mutually non-interacting feedforward pathways. This property is incompatible with the weight-RL mechanism of our model. For this reason, ReLU was not considered suitable for the present framework.

##### 4-6. Regularization

We used L2 regularization. Prior research has identified it as a strong candidate for enabling continual learning in sequential settings.[25] Beyond its well-known stabilizing effects, L2 regularization plays an additional role in our model by supporting the weight-RL mechanism, which must remain functional as a sequential adaptation channel throughout both the learning phase and the overtrained phase.
While stronger regularization generally promotes continual adaptation, excessively large coefficients interfere with the learning dynamics of the recurrent network. Because the goal of this study is not to fit the behavioral data, we did not perform exhaustive hyperparameter optimization. Nevertheless, the choice of regularization strength is practically important, and future research may benefit from a more systematic investigation of this parameter.

#### Supplementary Figures

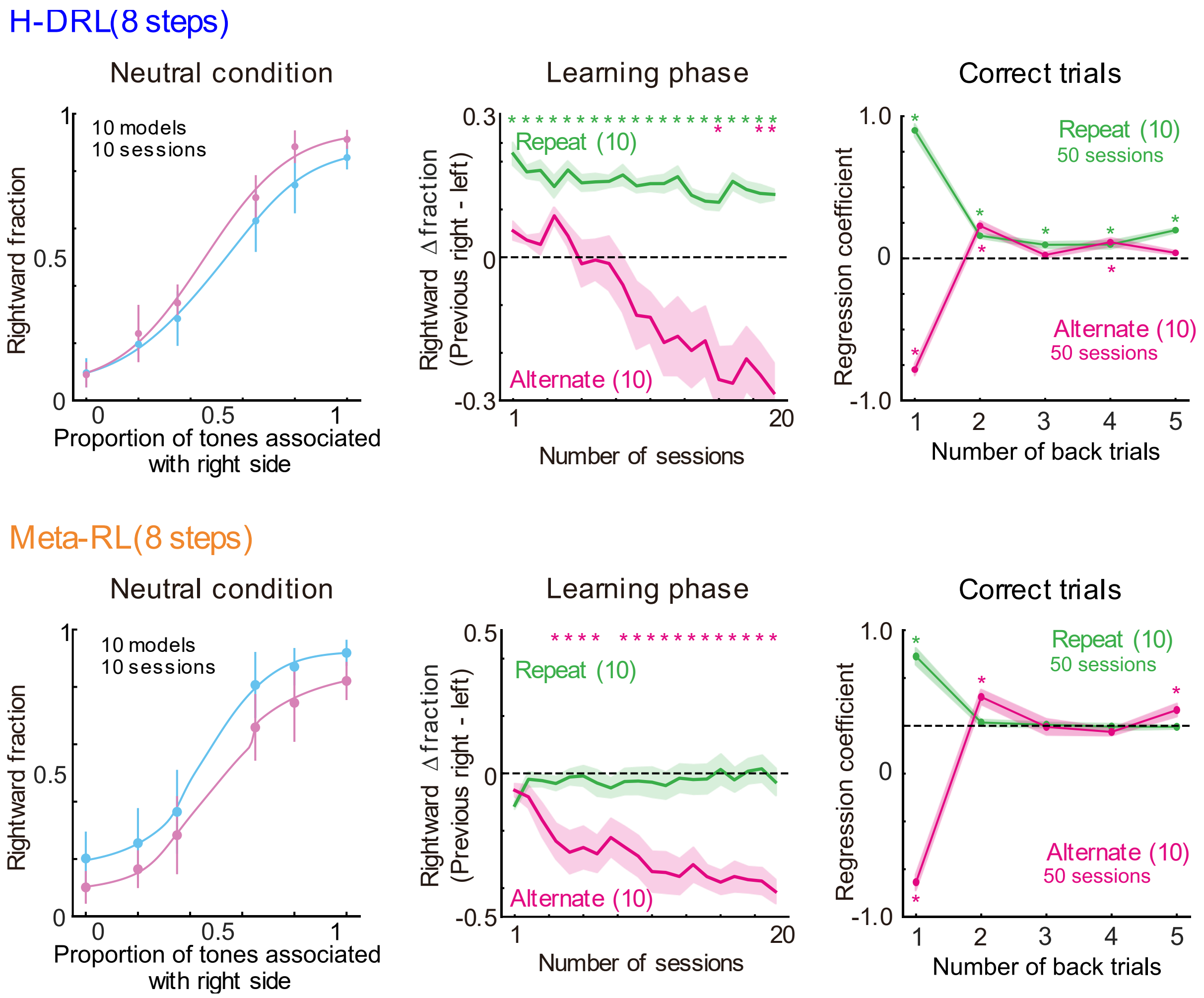


**Supplementary Fig. 1. Model behavior in the extended environment.**

(top) Analysis of H-DRL behavior in the case of training in the extended environment for **Fig. 5**. Data presentation is consistent with that in **Fig. 2**. (bottom) Analysis of Meta-RL behavior.

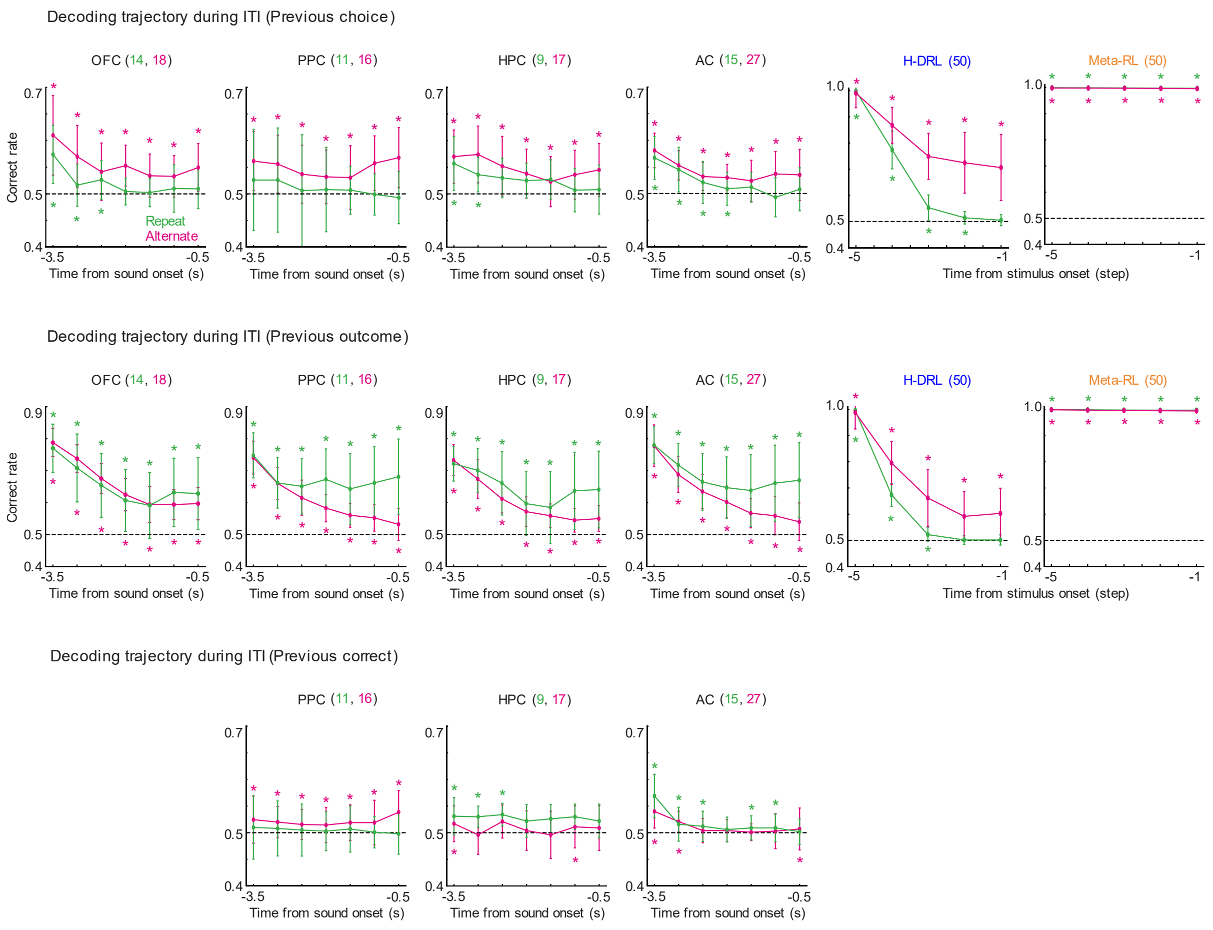


**Supplementary Fig. 2. Decoding the trajectory during ITI.**

Decoding of previous events from the activities of mouse neurons and model units. Data presentation is consistent with that in **Fig. 5d**.

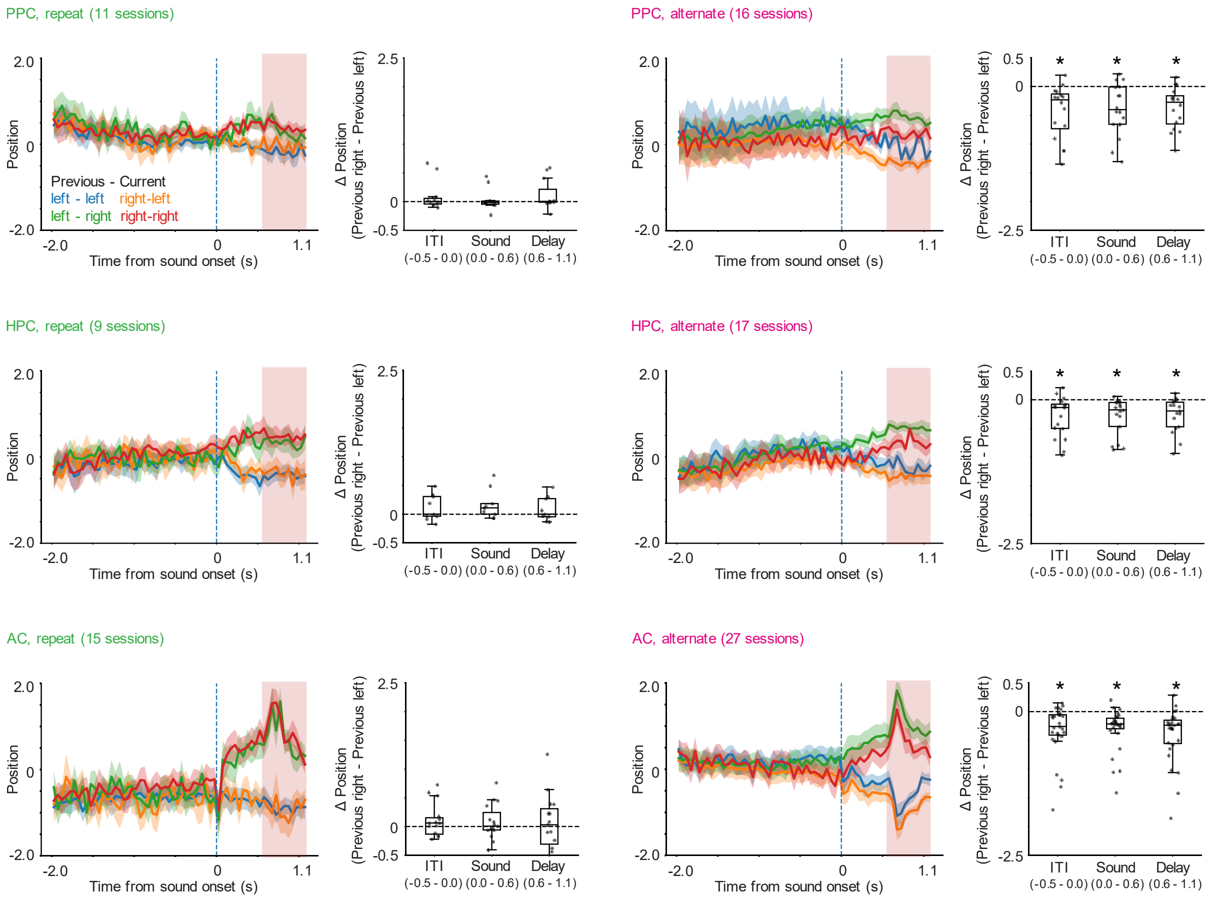


**Supplementary Fig. 3. Choice axis projection for PPC, HPC, and AC.**

Population activity of PPC, HPC, and AC neurons projected to a choice axis during the delay period. Data presentation is consistent with that in **Fig. 5e**.
